# GENEvaRX: A Novel AI-Driven Method and Web Tool Can Identify Critical Genes and Effective Drugs for Lichen Planus

**DOI:** 10.1101/2023.02.23.529678

**Authors:** Turki Turki, Y-h. Taguchi

## Abstract

Lichen planus (LP) is an autoimmune disorder diagnosed based on physical symptoms and lab tests. Examples of symptoms include flat bumps, and itchy and purplish skin, while lab tests include a shave biopsy of the lesion. When the pathology report shows consistency with LP and is negative for potential triggers for an allergy test and hepatitis C, a dermatologist typically prescribes corticosteroid in the form of pills or injection into the lesion to treat the symptoms. To understand the molecular mechanism of the disease and thereby overcome issues associated with disease treatment, there is a need to identify potential effective drugs, drug targets, and therapeutic targets associated the LP. Hence, we propose a novel computational framework based on new constrained optimization to support vector machines coupled with enrichment analysis. First, we downloaded three gene expression datasets (GSE63741, GSE193351, GSE52130) pertaining to healthy and LP patients from the gene expression omnibus (GEO) database. We then processed each dataset and entered it into our computational framework to select important genes. Finally, we performed enrichment analysis of selected genes, reporting the following results. Our methods outperformed baseline methods in terms of identifying disease and skin tissue. Moreover, we report 5 drugs (including, dexamethasone, retinoic acid, and quercetin), 45 unique genes (including PSMB8, KRT31, KRT16, KRT19, KRT17, COL3A1, LCE2D, LCE2A), and 23 unique TFs (including NFKB1, STAT1, STAT3) reportedly related to LP pathogenesis, treatments, and therapeutic targets. Our methods are publicly available in the GENEvaRX web server at https://aibio.shinyapps.io/GENEvaRX/.

## 1. Introduction

Lichen planus (LP) is a non-infectious autoimmune disorder in which the immune system attacks affect different areas, including the skin, mouth, and genitals, among other [1, 2]. Skin symptoms include itchy (or irritating), flat bumps, and purplish, characterized by more persistent symptoms depending on the affected area (e.g., genitals or mouth) [3]. Normally, a shave biopsy is performed by shaving the skin lesion and sending it to a pathology laboratory [4, 5]. Then, a dermatopathologist uses a microscope to inspect the sample skin for disease diagnosis. Examples of microscopic descriptions include skin associated with hypogranulosis, hyperkeratosis, and vacuolar basal change. This also involves specifying whether the skin dermis demonstrates band-like patterns of lymphocytic infiltration, pigment incontinence, and other info pertaining to stains [6–8]. Such descriptions provide a more accurate diagnosis if the skin lesion is related to LP. Other possible triggers of LP include hepatitis C and allergic reactions [9, 10]. When potential triggers are excluded, doctors prescribe treatments to manage LP and treat the symptoms. As the etiology for LP is unknown and the disease still has no cure [11, 12], researchers, mainly in biology, medicine, and computational biology domains, have attempted to describe the molecular mechanism of LP and gain biological insights, thereby speeding up the drug discovery process [13].

Vo et al. [14] utilized a bioinformatics approach to gain biological insights into oral lichen planus (OLP). The input datasets were downloaded from the gene expression omnibus (GEO) database according to the accession numbers GSE52130 and GSE38616. Both datasets have 14 uniformly distributed samples consisting of seven healthy controls and seven OLP patients. GSE52130 is related to the epithelium of OLP patients, while GSE38616 is related to the mucosa of OLP patients. A t-test was performed for each input dataset to identify differentially expressed genes (DEGs) with *p* < 0.05 and with a Benjamini-Hochberg FDR correction. A cluster analysis was performed on the GSE52130 dataset to remove outlier samples and thereby produce a partial dataset, for which a t-test was performed, to obtain 444 DEGs. Similarly, a cluster analysis was performed on GSE38616 to remove outlier samples, producing a partial dataset for which a t-test was performed, with an output of 348 DEGs. Then, taking the intersection of these DEGs (i.e., 444 and 348), 43 DEGs were obtained and input into DAVID (http://david.abcc.ncifcrf.gov) for gene ontology enrichment analysis. Results demonstrated that the 43 DEGs included genes related to wound healing (e.g., TNC, and TGFBI), hyperkeratosis (e.g., TMEM45A), and response to infection (TMEM173, LITAF, and RFTN1), among others related to OLP inflammation.

De Lanna et al. [15] performed a bioinformatics analysis to gain further biological insights regarding the malignant transformation of OLP by identifying shared molecular signatures, thereby distinguishing OLP from oral squamous cell carcinoma (OSCC). The input datasets are publicly available from the GEO (GSE52130, GSE56532, and GSE41613) and were related to OLP (and healthy controls), OSCC in advanced stages, and in stages I and II with the corresponding healthy controls. Analysis of co-expressed genes was performed using the Weighted Gene Coexpression Network Analysis (WGCNA) R package. Analysis of differentially expressed genes was performed on a given dataset using limma according to *p* < 0.05, with Benjamini-Hochberg correction. In the three datasets, 107 DEGs were identified from OLP and healthy controls, 331 DEGs for early stage (i.e., I and II) OSCC against healthy controls, and 282 DEGs for advanced stage OSCC against healthy controls. Among the datasets, 35 overlapping DEGs were considered as to be gene signatures of OLP and OSCC. Out of 35 genes, 18 are shared between OLP and OSCC. Immune-related genes, such as IFI6 and IFI27, were among the 35, as well as keratinocyte differentiation genes such as S100A12 and S100P. Enrichment analysis also identified drugs, including oncogenes such as PI3, SPRR1B and others related to the immune system that were also dysregulated in OLP, such as CXCL13, and HIF1A. Co-expressed genes were also identified, elucidating the shared molecular mechanism of OLP and OSCC.

Zhao et al. [16] aimed to inspect the therapeutic effects of quercetin on OLP. For quercetin, 75 targets were identified using two databases: TCMSP (http://tcmspw.com) and UniProt (https://www.uniprot.org). Additionally, 6110 differentially expressed genes of OLP were considered targets. The intersection of 75 and 6110 targets resulted in 66 common targets, which were entered into a network-based enrichment analysis, which functions follows: first, a PPI network was constructed by entering the 66 common targets into the STRING database (https://string-db.org/cgi/input.pl). A topological analysis of the PPI network was then performed using the cytoHubba tool, identifying 12 important targets (F3, VEGFA, TNF, SELE, TP53, JUN, CCL1, VCAM1, PLAT, IL1B, IL6, and IFNG). Second, the common 66 targets were input into a Gene Ontology (GO) enrichment analysis to retrieve biological processes, cell components, and molecular function terms. Kyoto Encyclopedia of Genes and Genomes (KEGG) enrichment analysis reported pathways based on the 66 common targets. Then, a drug–disease–pathway–target interaction network showing the relationship of quercetin and OLP was visualized via Cystoscape. Outcomes were then validated using in vivo experiments, in which higher concentrations of quercetin were associated with higher apoptosis rates of T lymphocytes. Results indicated that quercetin interfered with T helper cells (TH1/TH2) through the protein expression levels of cytokines (IL-6 and IFN-γ) to modulate immune response in OLP. Other recent studies have relied on bioinformatics-based approaches to gain biological insights contributing to understanding the molecular mechanism of LP [17–20].

Although current biological advances understanding of LP mainly relies on bioinformatics-based approaches with existing tools, as well as biological databases of limited knowledge, computational frameworks derived with the help of advanced concepts in artificial intelligence (AI) are needed to unlock tremendous biological knowledge and to lay the foundations for AI as a driving force. Such an existing tool can help in reducing the search space of identifying therapeutic targets, drug targets, and effective drugs, helping to advance drug discovery and disease understanding.

### Contributions

Our major contributions in this work can be summarized as follows:

1. We introduce a computational framework based on machine learning to extract more biological knowledge pertaining to LP. Our framework consists of three new variants of support vector machines (SVM) coupled with enrichment analysis tools, including Enrichr and Metascape [21, 22].
2. We downloaded three gene expression datasets pertaining to LP from the gene expression omnibus (GSE63741, GSE193351, and GSE52130). Then, we compared our framework against four baseline methods, including limma [23], significance analysis of microarrays (SAM), and *t*-test [24], and lasso for (1) the identification of LP skin disease; and (2) skin tissue identification.
3. Enrichment analysis results indicate the superiority of our methods when retrieving (1) related terms to LP with more expressed genes; and (2) terms related to skin tissue with more expressed genes induced by our methods.
4. Further enrichment analysis results report extensive biological insights as follows. In the first dataset, we identified drugs such as Alitretinoin, Vitinoin, and Dexamethasone, as well as their corresponding drug targets. We also identified proteasome (e.g., PSMB8), expressed genes related to Interferon alpha/beta signaling (e.g., IFL27 and IFITM1), and other keratin genes such as KRT17, all of which are related to LP T-cell mediated disease affecting the skin. Fifteen transcription factors were identified (e.g., NFKB1, RELA, and STAT1) that contribute to the LP inflammatory process. For the second dataset, expressed genes related to skin extracellular matrix (e.g., COL6A3, COL3A1) were identified, and interestingly, expressed genes related to ribosomes dysfunction (e.g., RPL35A, RPS27A, RPS14, RPSA), which contributes to autoimmune skin diseases, were reported. Additionally, drugs (Quercetin, Vitinoin, and Retinoic acid) and their corresponding drug targets were identified and four TFs (RELA, STAT3, TP53, STAT1) related to LP T-cell mediated autoimmune inflammation were reported. For the third dataset, three drugs (Retinoic acid, Vitinoin, and Alitretinoin) were reported with their associated drugs, as well as three expressed genes (PI3, LCE2A, LCE2D) related to the formation of the cornified envelope at the skin epidermis. Furthermore, LCP1 and ITGB2 expressed genes related to cytokine signaling in immune system were reported. Of the 11 TFs, STAT1, STAT3, RELA, and NFKB1 have been reported and involved in the immune attack to the skin.
5. Our tool can save time and cost in the drug discovery development process. For example, results show that our framework can successfully narrow the search for therapeutic targets, thereby reducing the time and drug choices in clinical trials and helping reduce the cost of drug discovery. This can also facilitate the identification of both drug targets and candidate drugs.
6. To reproduce the results of this study, we made an efficient implementation of our methods publicly available within the GENEvaRX web server at https://aibio.shinyapps.io/GENEvaRX/. We provide datasets in the Supplementary Datasets folder. We also provide a GENEvaRX_Screenshots.docx file within the Supplementary Information to demonstrate the use of the web tool.

### Organization

The rest of this paper is organized as follows: In Section 2, we provide a description of gene expression datasets employed in this study, followed by and introduction to our computational framework. In Section 3, we conduct an experimental study, evaluating and reporting the performance of our methods against existing baselines. A discussion is provided in Section 4. Finally, we conclude our work and point to future directions for research in Section 5.

## 2. Materials and Methods

### 2.1 Gene Expression Profiles

We downloaded three gene expression datasets with the following accession numbers from the GEO database [25].

#### 2.1.1 GSE63741: Dataset1

This gene expression dataset consisted of 60 samples and 1542 expression values [26]. Therefore, we had a 60 × 1543 matrix, including a column vector for the sample type. The sample type distribution was unified, in which 30 samples were designated as “Control” while the remaining samples were from “Lichen Planus” patients. The gene expression dataset was measured and performed according to the microarray platform PIQOR (TM) Skin 2.0 Microarray, human, antisense (591). We refer to this dataset as Dataset1.

#### 2.1.2 GSE193351: Dataset2

We retrieved the whole transcriptomic dataset comprised of 47 samples and 64253 expression values [25]. Then, we downloaded a list of approved protein-coding genes (PCGs) by domain experts from BioMart at HUGO Gene Nomenclature Committee (HGNC) at https://biomart.genenames.org/ [27, 28]. We selected 5916 PCGs from the 64253 genes. Thus, we ended up with a dataset encoded as a 47 × 5917 matrix, including the sample type as a column vector. The sample type distribution was not unified, consisting of 10 healthy controls and 37 LP patients. The dataset was then measured and run using the high throughput sequencing platform Illumina HiSeq 4000 (Homo sapiens). We refer to this dataset as Dataset2.

#### 2.1.3 GSE52130: Dataset3

The third gene expression dataset included 23 samples and 43961 expression values [25]. We selected 5355 approved PCGs by domain experts from BioMart at HUGO Gene Nomenclature Committee (HGNC) at https://biomart.genenames.org/ [27, 28]. The sample distribution of 14 samples was uniformed, consisting of 7 samples for each class. Therefore, we had a dataset encoded as a 14 × 5356 matrix, including the sample type as a column vector. The sample type distribution was comprised of 14 samples (distributed as 7 healthy controls, 7 OLP patients) and 9 samples (distributed as 5 genital lichen planus and 4 healthy controls). The gene expression data was measured and processed according to the microarray platform Illumina HumanHT-12 V4.0 expression beadchip. We refer to this dataset in the remaining sections as Dataset3.

### 2.2 Computational Framework

Figure 1 demonstrates the performed steps of the presented computational framework. Given a gene expression dataset as {(x_1_,y_1_),…, (x_m_,y_m)_}, where x_i_ designates the i*th* sample and y_i_ is the corresponding sample type. In this study, we considered y_i_ in the binary case ({Control, Lichen Planus}). We encoded all samples x*_i_* (for *i =* 1..*m*) as an *m* × *n* matrix, where *m* and *n* are the number of samples and genes, respectively. We also had a 1 × *m* column vector, encoding all sample types y*_i_* (for *i =* 1..*m*) as. We aimed to identify *p* important genes out of the *n* genes, where *p* « *n*, and these *p* genes are expressed in each sample. Therefore, we presented new sparse variants of the SVM, working as follows: in Equation 1, we sought to find arguments (i.e., w = [*w*_1_…*w*_n_] and bias *b* ∈ R) minimizing the objective function subject to shown constraints. Each sample was encoded by expression values of *n* genes. As a result, weights in w corresponded to the gene importance among all samples. A higher weight *w*_i_ refers to greater importance of the *i*th gene.

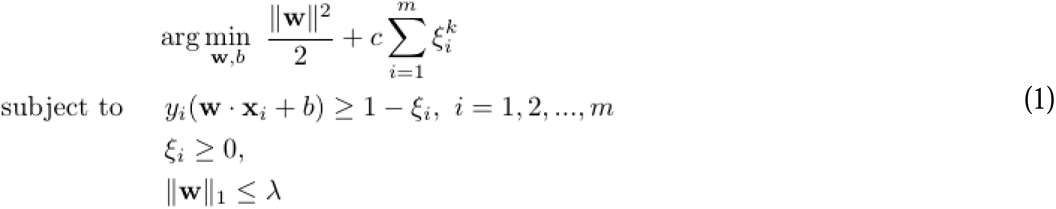

**Figure 1:**
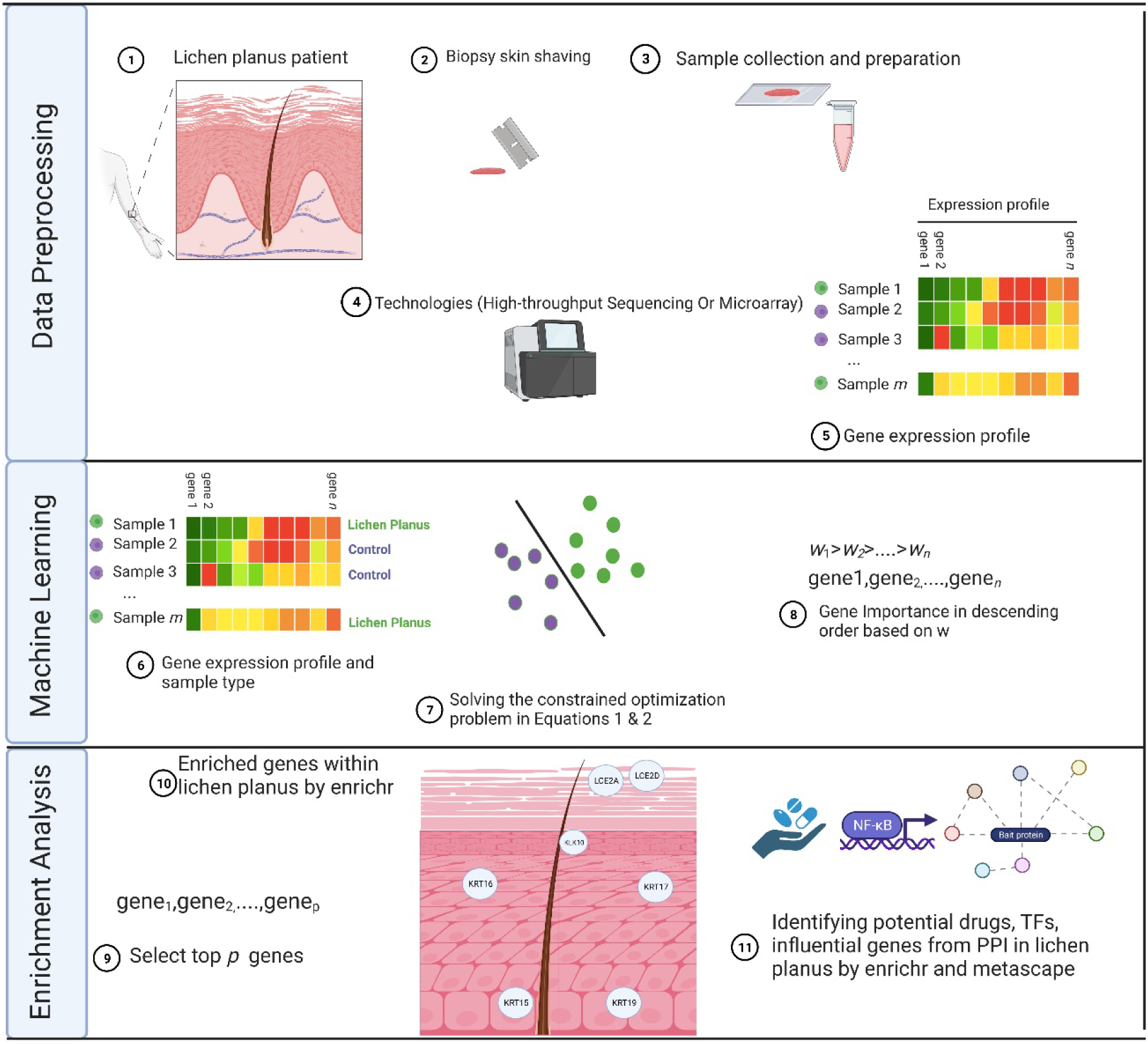
Flowchart of the presented AI-driven computational framework identifying transcription factors, drugs and important genes expressed in LP. Figure created with BioRender.com.

In Equation 1, the constrained optimization problem is different from the soft margin SVM in [22], as it includes additional constraint IIW II_1_ ≤ λ, shrinking weights for genes with less prediction power closer to zero. That leads to increased sparsity for weight vector 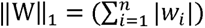 is the *L*_1_-norm, λ ∈ ℝ. Equation 2 shows a new variant of the constrained optimization problem by including IIW II_1_ ≤ λ as an additional constraint. Therefore, when |*v*| > *r*, then Equation 2 becomes the formula shown in Equation 3. Otherwise, the constrained optimization problem becomes as shown in Equation 4. For a given input dataset, we had different w and *b* as a result of the different optimization problems. We utilized CVXR in R to find the weight vector w and bias *b,* minimizing the constrained optimization problem [29].

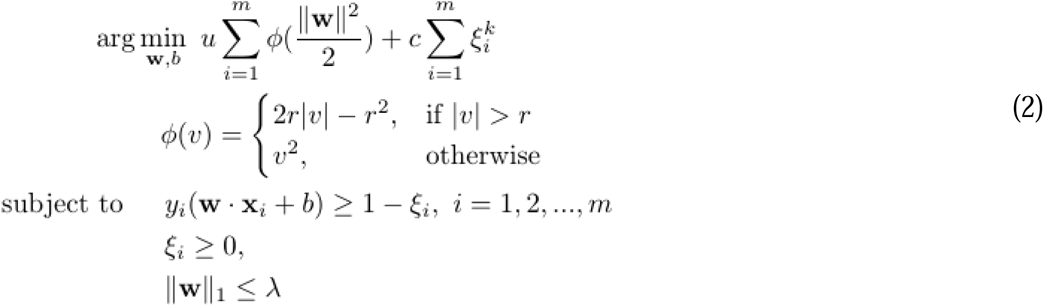

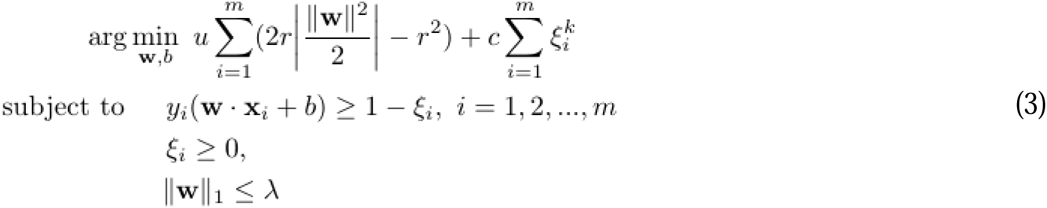

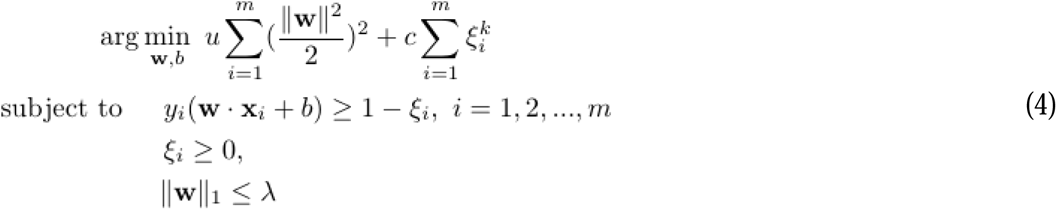

### 3 Experiments and Results

#### 3.1 Experimental Methodology

We assessed our presented methods (i.e., ssvm1, ssvm2 and ssvm3) against several baseline methods, including limma [23], SAM, *t*-test, and lasso [24]. All seven approaches received labeled examples as an input to the computational method. For our methods (i.e., ssvm1, ssvm2, and ssvm3), because these SVM-based models are defined as the sign(w.x + *b*) (i.e., sign of weighted genes w.x plus the bias *b*), the top *p* important genes are selected based on the corresponding top *p* values in weight vector, as those top *p* values in w (i.e., w = [*w*_1_…*w_p_*]) refer to the importance of selected genes. For lasso, the model is expressed as *β*_0_ + *β*x, where top *p* genes are associated with top *p* coefficients in *β*. Hence, we selected top *p* genes based on top *p* coefficients in /3. For baseline methods (i.e., limma, SAM, *t*-test), we selected genes based on significance according to adjusted *p*-values < 0.01. In Table 1, we list all methods employed in this study.

**Table 1:** List of employed computational methods in this study.

| Abbreviation | Method |
| --- | --- |
| ssvm1 | Sparse svm 1 (proposed) |
| ssvm2 | Sparse svm 2 (proposed) |
| ssvm3 | Sparse svm 3 (proposed) |
| limma | Linear models for microarray data (baseline) |
| sam | Significance analysis of microarrays (baseline) |
| $t$-test | Student's $t$-test (baseline) |
| lasso | Least absolute shrinkage and selection operator (baseline) |

We performed a biological enrichment analysis evaluating the performance results by uploading genes produced via each method to Enrichr (https://maayanlab.cloud/Enrichr/) [30] and Metascape (https://metascape.org/gp/index.html) [31]. Assuming that the retrieved term was about a disease, then more genes associated with a disease demonstrated better performance when the disease was related to the studied disease type. The same concept holds true when a term is a tissue (i.e., composed of cells with the same function). Another performance indicator was the ranking, where a lower ranking of the disease (e.g., lichen planus) showed the better performance results. We also evaluated our models against lasso using the supervised learning setting, reporting performance measures, such as area under curve (AUC), followed by plots related to statistical significance of results and gene importance. In this study, we used R to conduct the experiments [32]. In particular, we employed the CVXR package in R to solve optimization problems [29]. We also performed SAM utilizing the siggenes package [33] and we employed the limma package in R, using the lmFit and eBayes functions for model fitting [23]. For the t-test, we used the t.test function in the R stats package [32]. For lasso, the glmnet package [34] was used to select genes associated with non-zero coefficients, as these refer to the important genes. For the three baseline methods (i.e., SAM, limma, and *t*-test), we employed the p.adjust function coupled with the “BH” option to compute adjusted *p*-values, setting a threshold of *p* < 0.01 as in [22, 35].

### 3.2 Results

#### 3.2.1 Dataset1

For each method, we show retrieved terms provided by Enrichr in Tables 2 and 3. These terms in Table 2 show the diseases associated with provided input list of genes. Related terms with low-ranking values and a large number of expressed genes demonstrated the superiority of the computational method. Table 2 shows that ssvm3 performed better than all other methods. Notably, lichen planus is ranked 27^th^, with 4 expressed genes (CXCL9, MMP9, S100A9, S100A8) out of 38. Moreover, autoimmune diseases is ranked 4^th^, with 20 expressed genes (CD52, RARRES3, STAT1, TRAC, ITGB2, GSTT1, ISG15, MMP9, PSMB10, PSMB8, TNFSF13B, PSMB9, KRT19, SOCS1, GAL, LCK, IRF1, CD47, S100A9, S100A8) out of 1060. The ssvm1 and ssvm2 methods outperformed ssvm3 and other methods for dermatologic disorders term, ranking 9th with 12 commonly expressed genes (MMP12, SOCS1, KRT17, RARRES3, ITGB2, TNC, CCL19, MMP9, MS4A1, CST6, S100A8, CCL27). Table 3 shows retrieved terms related to tissues of interest. It is clear that ssvm3 achieved the best results regarding skin tissue, ranking 3rd with 20 expressed genes (SERPINA12, CXCL9, MAOA, TNC, KRT7, CST6, MMP12, FADS2, KRT19, GAL, KRT17, COL4A1, KRT16, KRT15, SFN, THRSP, CCL19, S100A9, S100A8, CCL27) within the skin tissue comprised of cells with similar function. Tied for the second-best method regarding skin tissue, ssvm2 and ssvm3 ranked 6^th^, with 19 expressed genes. When compared to baseline methods, these results demonstrate the superiority of ssvm3, followed by ssvm1 and ssvm2. Although skin tissue was ranked the 6th by SAM, 13 genes were expressed, fewer than those of ssvm2 and ssvm3. Figure 2 provides useful information pertaining to the intersection of provided genes according to each method. It can be seen that ssvm1, ssvm2, and ssvm3 share 70 genes, while they differ in 30 genes. This implies that each method generated different list of genes. For example, limma has 79 different genes from all other methods. The same holds for SAM. The t-test showed 75 different genes from all other methods. Additionally, lasso produced 12 genes in which 7 were unique to lasso (i.e., no other method produced these genes). These visualized results act as a demonstration of performance differences between methods, as each produced a different gene list. Regarding Dataset1, we provide genes produced by all computational methods in Supplementary Data Sheet 1. In Supplementary Tables 1_A and 1_B, we provide enrichment analysis results of all methods pertaining to DisGeNET and ARCHS4 Tissues, respectively.

**Table 2:**
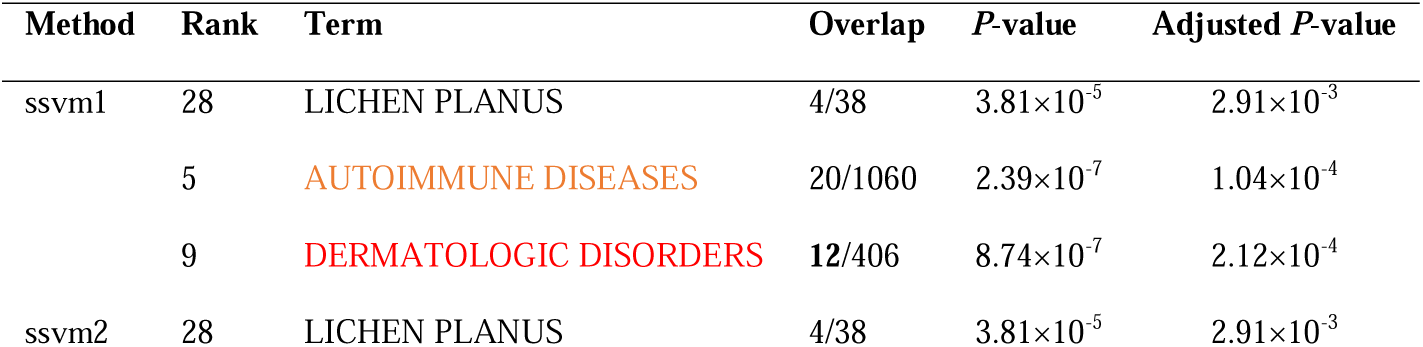
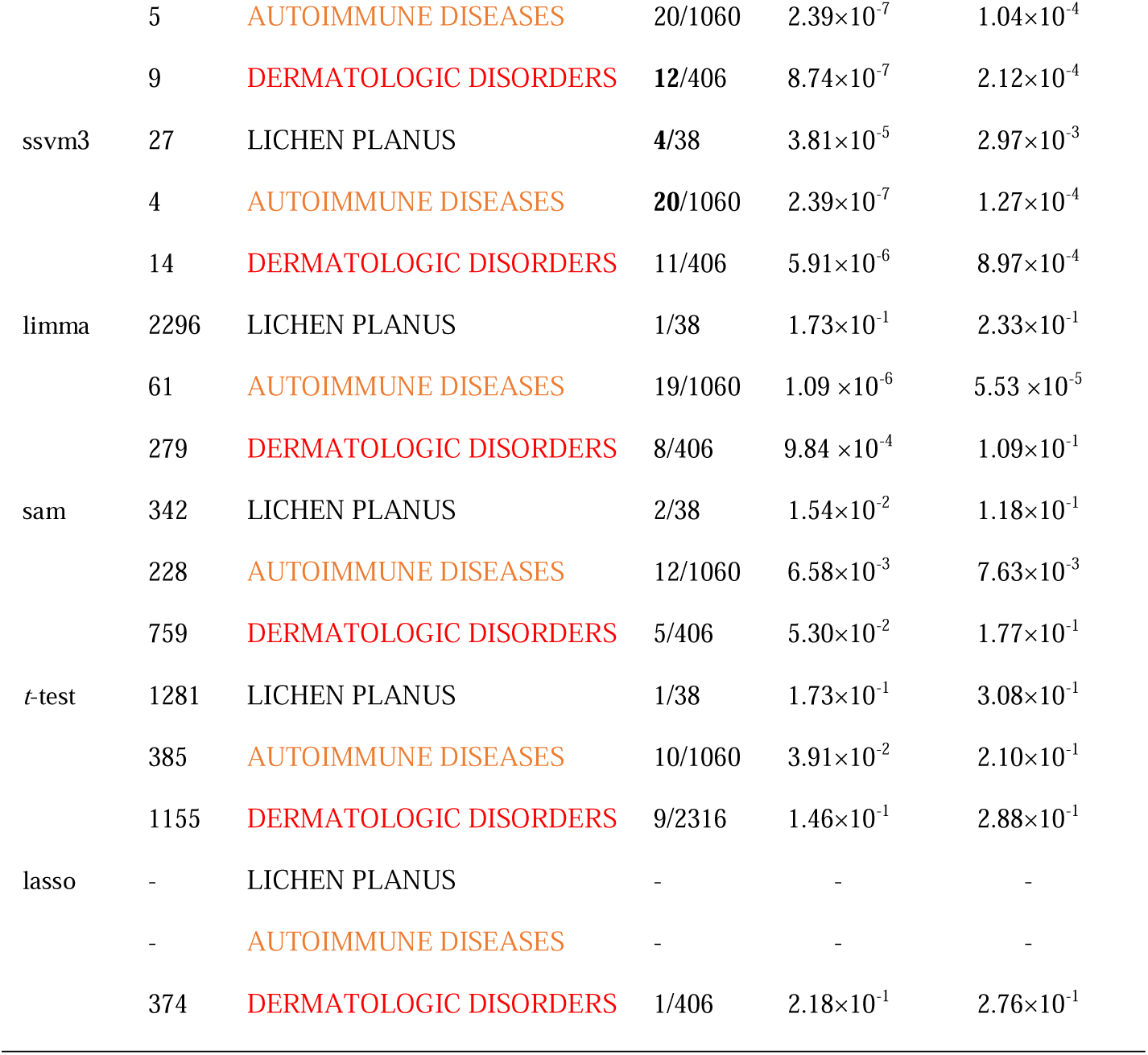
Retrieved enriched terms within “DisGeNET” via Enrichr according to uploaded genes via methods using Dataset1, showing association between human genes and diseases. Rank column shows the order of terms when retrieved. Overlap displays a ratio-like format, corresponding to the number of uploaded genes overlapping to those in the terms. The best result of each method is shown in bold.

**Table 3:** Retrieved enriched terms within “ARCHS4 Tissues” via Enrichr according to uploaded genes via methods using Dataset1, showing genes expressed within tissue types. Rank column shows the order of terms when retrieved. Overlap displays a ratio-like format, corresponding to the number of uploaded genes overlapping to those in the terms. The best result of each method is shown in bold.

| Method | Rank | Term | Overlap | P-value | Adjusted P-value |
| --- | --- | --- | --- | --- | --- |
| ssvm1 | 6 | SKIN (BULK TISSUE) | 19/2316 | $1.99 \times 10^{-2}$ | $3.59 \times 10^{-1}$ |
| ssvm2 | 6 | SKIN (BULK TISSUE) | 19/2316 | $1.99 \times 10^{-2}$ | $3.59 \times 10^{-1}$ |
| ssvm3 | **3** | SKIN (BULK TISSUE) | **20**/2316 | $1.00 \times 10^{-2}$ | $3.60 \times 10^{-1}$ |
| limma | 13 | SKIN (BULK TISSUE) | 20/2316 | $1.00 \times 10^{-2}$ | $7.72 \times 10^{-2}$ |
| sam | 6 | SKIN (BULK TISSUE) | 13/2316 | $3.72 \times 10^{-1}$ | $9.99 \times 10^{-1}$ |
| *t*-test | 9 | SKIN (BULK TISSUE) | 12/2316 | $4.93 \times 10^{-1}$ | $9.99 \times 10^{-1}$ |
| lasso | 9 | SKIN (BULK TISSUE) | 2/2316 | $4.12 \times 10^{-1}$ | $7.71 \times 10^{-1}$ |

**Figure 2:**
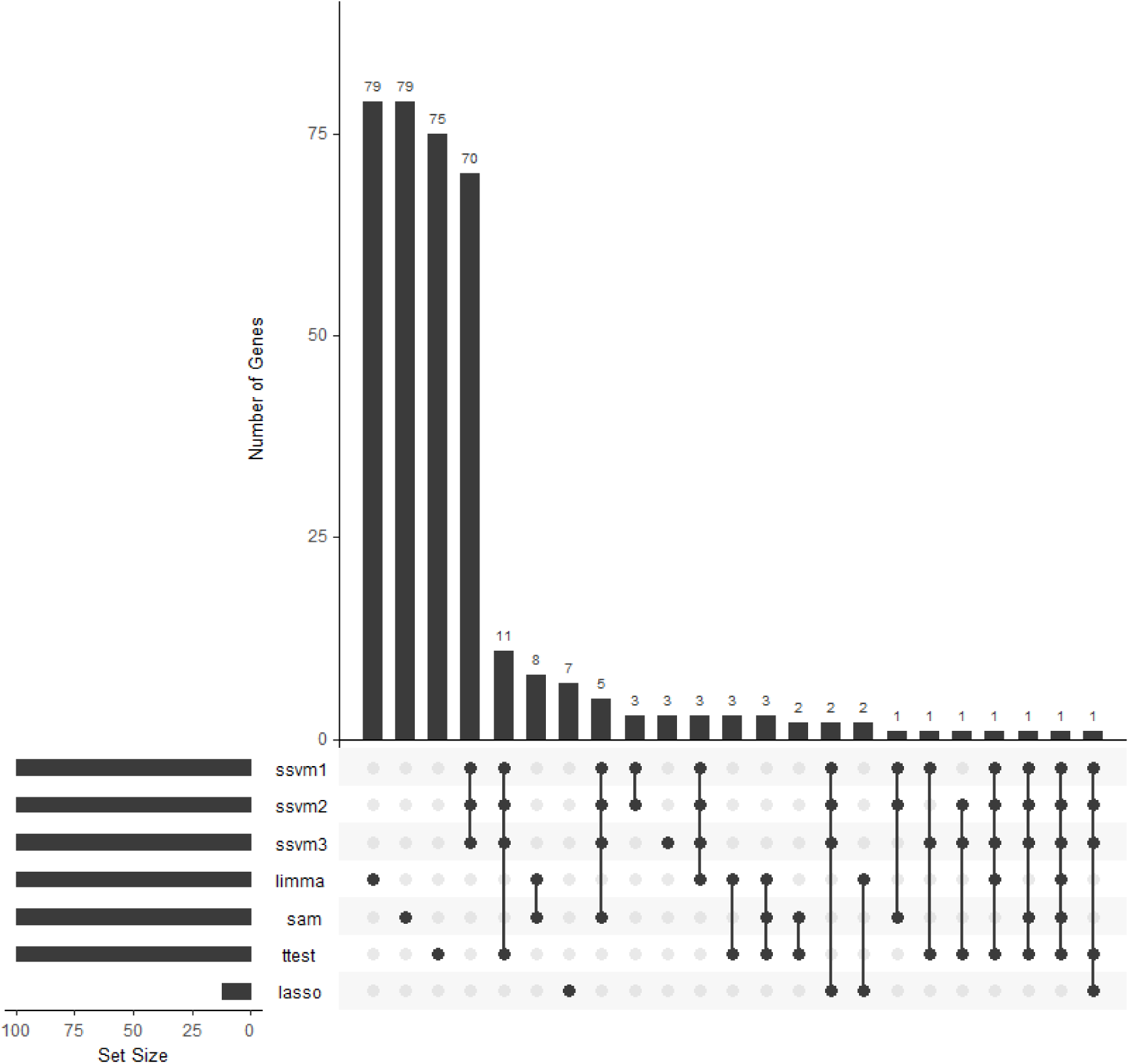
Visualization of notable sets pertaining to each gene list provided by each method when Dataset1 was used.

Because ssvm3 produced good results in terms of retrieving the diseases and tissues pertaining to the gene expression profiles in Dataset1, we retrieved drugs and associated drug targets (genes) from the drug signatures database (DSigDB) and then uploaded 100 genes of ssvm3 to Metascape for further biological analysis. Table 4 shows retrieved terms pertaining to drugs associated with provided genes. The first term is for the drug Alitretinoin, which has been shown to be clinically effective in the treatment of LP cases [36, 37]. The 2nd-ranked drug was Vitinoin, the generic name of which is tretinoin, which has been included in clinical trials and has demonstrated potential improvements in the clinical treatments pertaining to LP [38, 39]. Dexamethasone is a steroid-based drug ranked 16th and has demonstrated good clinical outcomes associated with OLP [40–43]. In Supplementary Table 1_C, we provide enrichment analysis results pertaining to drug terms within DSigDB. In Figure 3, we show three highly connected subgraphs obtained from the PPI network, including the following 14 genes: PSME2, PSMB10, PSMB9, KRT31, KRT16, KRT19, KRT17, KRT15, BST2, STAT1, IFI27, IFITM1, PSMB8, and IRF1.

**Table 4:** Retrieved enriched terms from “DSigDB” via Enrichr according to uploaded genes using Dataset1, showing genes (column: Genes) associated with drugs (column: Term). Rank column shows the order of terms when retrieved. NO. shows the number of genes.

| Rank | Term | NO. | Genes |
| --- | --- | --- | --- |
| 1 | Alitretinoin | 11 | KRT19;SOCS1;KRT17;RARRES3;STAT1;IRF1;<br>KRT15;ITGB2; KRT7;S100A9;S100A8 |
| 2 | Vitinoin | 19 | CD52;IFITM1;ITGB2;LAPTM5;KRT7;MMP9;FADS2;<br>KRT19;SOCS1;GAL;KRT17;BASP1;LCK;WIF1;KRT16;IRF1;<br>KRT15;S100A9;S100A8 |
| 16 | Dexamethasone | 9 | MMP12;SOCS1;MAOA;WIF1;TNC;S100A9;MMP9;CST6;S100A8 |

**Figure 3:**
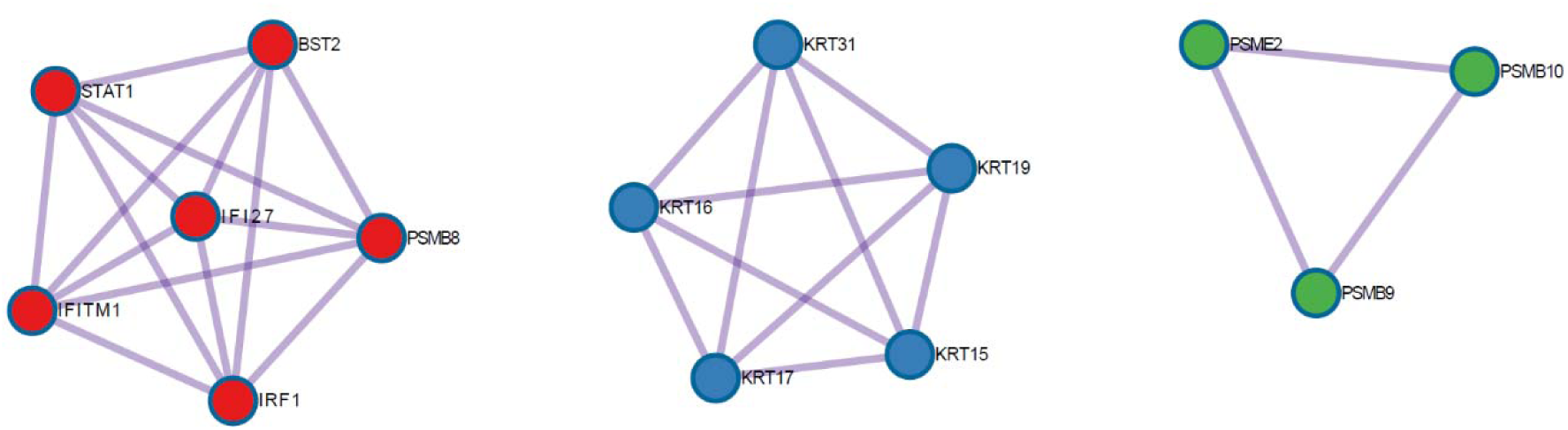
Three highly interconnected clusters in a protein-protein interaction based on 100 genes from ssvm3 provided to metascape enrichment analysis.

Some of these identified genes have played a key role in understanding the working mechanism of LP. Proteasome (e.g., PSMB9, PSMB8, PSMB10, PSME2) where PSMB8 dysfunction reported to play a key role in the induction od Type I interferon (IFN_α_,IFN_β_) [44]. We report expressed genes related to Interferon alpha/beta signalling (i.e., STAT1, BST2, IFL27, IFITM1, PSMB8, IRF1), as well as keratin genes related to intermediate filament organization (e.g., KRT16, KRT15, KRT17, KRT19, and KRT31). Regarding the persistence of wounds in skin autoimmune disease, it has been reported that Type I interferon, in addition to its role in dendritic cells activation and then T cells activation, can cause keratinocytes to attack [45]. These results demonstrate how T-cells attack the epidermis of the skin (Figure 4) [46], contributing to LP pathogenesis. It is worth noting that KRT17 has been reported to be a potential therapeutic targets for OLP [15]. Keratin 19 (KRT19) has been found to be expressed in OLP [47, 48]. KRT16 has been reported to be associated with LP in the wound repair process [14] and IFI27 has been reported to be overexpressed in OLP [15]. IRF1 and IFITM1 have been induced in LP, demonstrating their involvement in the skin injury process [49]. These 14 expressed genes can be considered as potential therapeutic targets for LP treatment. Moreover, they contribute to a greater understanding of LP pathogenesis.

**Figure 4:**
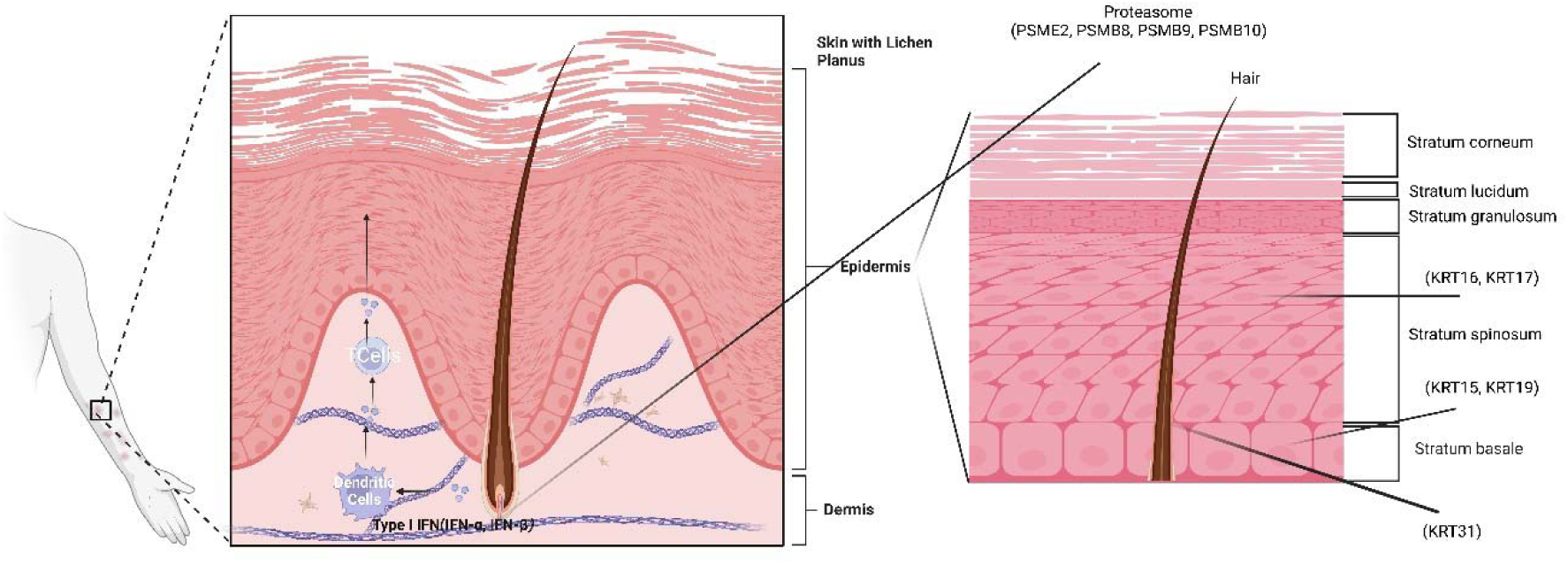
Skin of a patient, with lichen planus affecting the arm area. Critical expressed genes obtained from Figure 3 are shown in the skin epidermis and dermis. Figure created with BioRender.com.

We identified 15 transcription factors: RELA, NFKB1, SP1, STAT1, ETS1, STAT3, STAT2, JUN, HIF1A, IRF1, SRF, ETS2, CIITA, SMAD3, and SPI1. It has been reported that NFKB1 and RELA are among the crucial transcription factors in LP [50]. STAT1 has been reported to play a key role in the immune response in LP [19]. OLP has been reported to be associated with increased expression of SMAD3 [51]. These 14 identified genes and 15 transcription factors may prove useful as therapeutic targets and for understanding the molecular mechanism of LP.

#### 3.2.2 Dataset2

Table 5 reports terms (i.e., diseases) associated with selected genes while Table 6 reports terms (i.e., tissue type) according to expressed genes after uploading genes obtained via each method. It can be seen in Table 5 that our methods (ssvm1, ssvm2, and ssvm3) outperformed baseline methods. Specifically, skin diseases-genetic ranked 10th with 7 commonly expressed genes (DSP, FLG2, COL7A1, SPINK5, PKP1, ADAR, ITGA6) out of 45. Autoimmune diseases ranked 421st with 13 commonly expressed genes (FLG, SPARC, DST, STAT1, ADAR, ENO1, HMGCR, TTN, PTPRC, SOAT1, LMNA, FGFR3, TP63) out of 1060 genes. The dermatologic disorders term was ranked 13th with 14 commonly expressed genes (FLG, DSP, DST, SPINK5, DHCR24, ADAR, SFPQ, DES, COL7A1, LMNA, FGFR3, TP63, SPRR1B, IVL) out of 406. Baseline methods demonstrated inferior results, as retrieved terms ranked worse and exhibited fewer numbers of expressed genes within the retrieved terms. Moreover, no baseline method was able to identify all terms (e.g., skin diseases-genetic, autoimmune diseases, dermatologic disorders).

**Table 5:** Retrieved enriched terms within “DisGeNET” via Enrichr according to uploaded genes via methods using Dataset2, showing association between human genes and diseases. Rank column shows the order of terms when retrieved. Overlap displays a ratio-like format, corresponding to the number of uploaded genes overlapping to those in the terms. The best result of each method is shown in bold.

| Method | Rank | Term | Overlap | *P*-value | Adjusted *P*-value |
| --- | --- | --- | --- | --- | --- |
| ssvm1 | 10 | SKIN DISEASES, GENETIC | **7/45** | $2.45 \times 10^{-9}$ | $7.93 \times 10^{-7}$ |
| | 421 | AUTOIMMUNE DISEASES | **13/1060** | $2.37 \times 10^{-3}$ | $1.82 \times 10^{-2}$ |
| | 13 | DERMATOLOGIC DISORDERS | **14/406** | $1.46 \times 10^{-8}$ | $3.57 \times 10^{-6}$ |
| ssvm2 | 10 | SKIN DISEASES, GENETIC | **7/45** | $2.45 \times 10^{-9}$ | $7.93 \times 10^{-7}$ |
| | 421 | AUTOIMMUNE DISEASES | **13/1060** | $2.37 \times 10^{-3}$ | $1.82 \times 10^{-2}$ |
| | 13 | DERMATOLOGIC DISORDERS | **14/406** | $1.46 \times 10^{-8}$ | $3.57 \times 10^{-6}$ |
| ssvm3 | 10 | SKIN DISEASES, GENETIC | **7/45** | $2.45 \times 10^{-9}$ | $7.93 \times 10^{-7}$ |
| | 421 | AUTOIMMUNE DISEASES | **13**/1060 | $2.37 \times 10^{-3}$ | $1.82 \times 10^{-2}$ |
| | 13 | DERMATOLOGIC DISORDERS | **14**/406 | $1.46 \times 10^{-8}$ | $3.57 \times 10^{-6}$ |
| limma | - | SKIN DISEASES, GENETIC | - | - | - |
| | 944 | AUTOIMMUNE DISEASES | 6/1060 | $4.38 \times 10^{-1}$ | $6.58 \times 10^{-1}$ |
| | 795 | DERMATOLOGIC DISORDERS | 3/406 | $3.31 \times 10^{-1}$ | $5.92 \times 10^{-1}$ |
| sam | - | SKIN DISEASES, GENETIC | - | - | - |
|  | - | AUTOIMMUNE DISEASES | - | - | - |
|  | - | DERMATOLOGIC DISORDERS | - | - | - |
| t-test | - | SKIN DISEASES, GENETIC | - | - | - |
| | 578 | AUTOIMMUNE DISEASES | 1/1060 | $6.81 \times 10^{-1}$ | $7.00 \times 10^{-1}$ |
|  | - | DERMATOLOGIC DISORDERS | - | - | - |
| lasso | - | SKIN DISEASES, GENETIC | - | - | - |
| | 516 | AUTOIMMUNE DISEASES | 1/1060 | $6.03 \times 10^{-1}$ | $6.40 \times 10^{-1}$ |
| | 445 | DERMATOLOGIC DISORDERS | 1/406 | $2.94 \times 10^{-1}$ | $3.61 \times 10^{-1}$ |

**Table 6:**
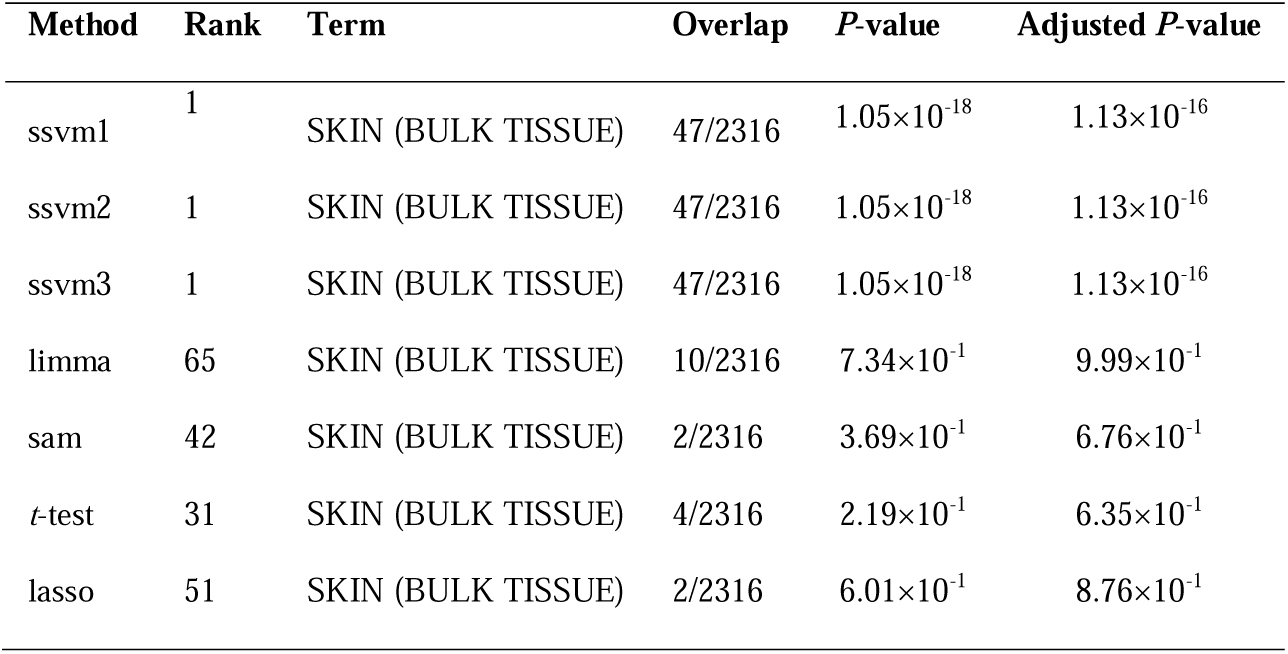
Retrieved enriched terms within “ARCHS4 Tissues” via Enrichr according to uploaded genes via methods using Dataset2, showing genes expressed within tissue types. Rank column shows the order of terms when retrieved. Overlap displays a ratio-like format, corresponding to the number of uploaded genes overlapping to those in the terms. The best result of each method is shown in bold.

| Method | Rank | Term | Overlap | P-value | Adjusted P-value |
| --- | --- | --- | --- | --- | --- |
| ssvm1 | 1 | SKIN (BULK TISSUE) | 47/2316 | $1.05 \times 10^{-18}$ | $1.13 \times 10^{-16}$ |
| ssvm2 | 1 | SKIN (BULK TISSUE) | 47/2316 | $1.05 \times 10^{-18}$ | $1.13 \times 10^{-16}$ |
| ssvm3 | 1 | SKIN (BULK TISSUE) | 47/2316 | $1.05 \times 10^{-18}$ | $1.13 \times 10^{-16}$ |
| limma | 65 | SKIN (BULK TISSUE) | 10/2316 | $7.34 \times 10^{-1}$ | $9.99 \times 10^{-1}$ |
| sam | 42 | SKIN (BULK TISSUE) | 2/2316 | $3.69 \times 10^{-1}$ | $6.76 \times 10^{-1}$ |
| t-test | 31 | SKIN (BULK TISSUE) | 4/2316 | $2.19 \times 10^{-1}$ | $6.35 \times 10^{-1}$ |
| lasso | 51 | SKIN (BULK TISSUE) | 2/2316 | $6.01 \times 10^{-1}$ | $8.76 \times 10^{-1}$ |

For example, SAM had no related terms. Hence, results were designated as “-”. The same held true when a baseline method didn’t retrieve the term. In Table 6, it is evident that our methods (i.e., ssvm1, ssvm2, and ssvm3) retrieve the related term (i.e., SKIN (BULK TISSUE)), ranking 1st with 47 expressed genes out of 2316. Baseline methods had worse ranking results. These results in Tables 5 and 6 demonstrate the superiority of our methods for disease identification as well as identification of tissues related to the studied problem. Figure 6 provided useful information pertaining to intersecting genes of each provided method. Our methods shared 98 genes in common while they differed in only 2 genes. Limma had 98 different genes from all other methods. The t-test and lasso methods differed by 16 and 9 genes, respectively. Our methods and limma shared 1 gene in common. The same was true for our methods relative to the t-test. These visualized results demonstrate that our methods differ from existing methods, and therefore results were different in Tables 5 and 6. In Supplementary Data Sheet 2, we provide a list of genes produced by each method used in enrichment analysis. Moreover, we provide enrichment analysis results pertaining to Tables 5 and 6 in Supplementary Table 2_A and Table 2_B, respectively.

**Figure 5:**
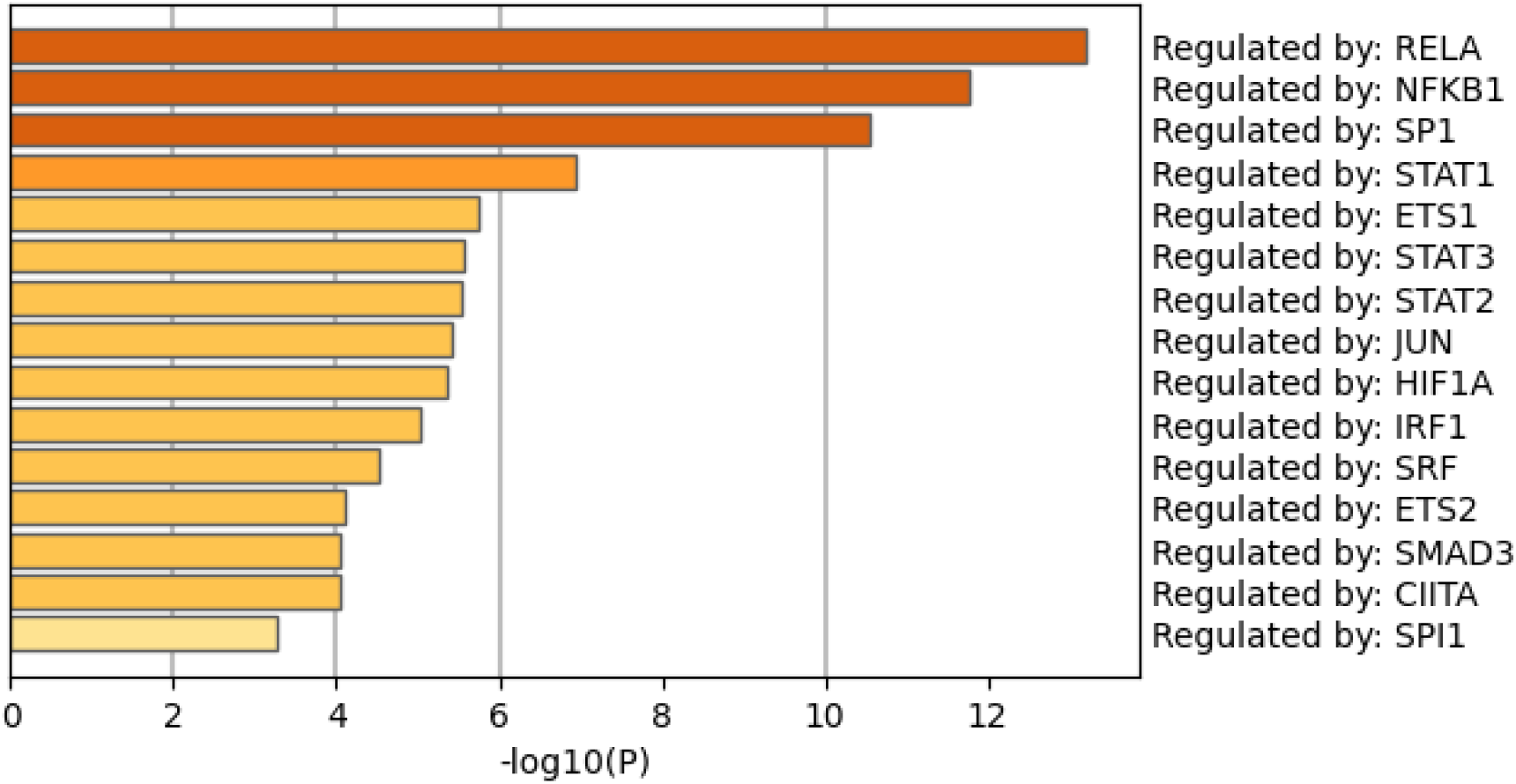
Fifteen transcription factors provided by Metascape according to the list of 100 input genes from ssvm3.

**Figure 6:**
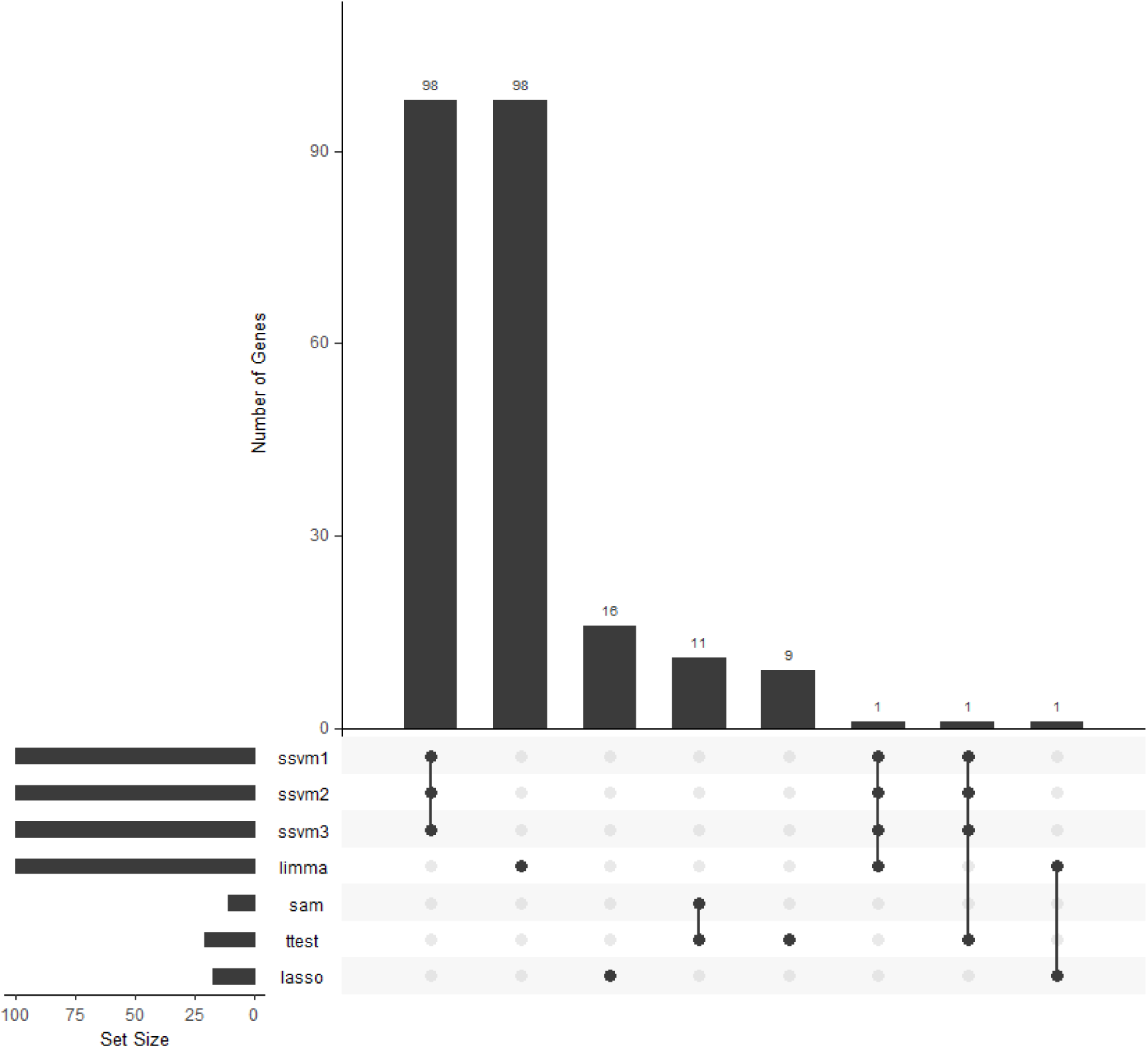
Visualization of intersecting sets pertaining to each gene list provided by each method when Dataset2 was used.

As our methods performed better than baseline methods and shared 98 genes, we selected ssvm3 to gain further biological insights within the molecular mechanism of LP. We retrieved drugs associated with genes produced via ssvm3 from Enrichr, followed by uploading the produced 100 genes into Metascape. Table 7 shows the drug terms and associated genes obtained from Enrichr. Retinoic acid has been reported to be effective as a topical treatment in OLP [52, 53]. Oral treatment of tretinoin (a generic name of Vitinoin) has been reported to be effective in treating LP [36]. In Supplementary Table 2_C, we provide enrichment analysis results pertaining to drug terms within DSigDB

**Table 7:** Retrieved enriched terms within “DSigDB” via Enrichr according to uploaded genes using Dataset2, showing genes (column: Genes) associated with drugs (column: Term). Rank column shows the order of terms when retrieved. NO. shows the number of genes.

| Rank | Term | # | Genes |
| --- | --- | --- | --- |
| 8 | Retinoic acid | 43 | EIF4A2;NOTCH2;FLG;MCTP1;HSP90AB1;EPAS1;SYNPO2;ITPR3;<br>ENO1;ACTG2;PTPRF;MYLK;RPS14;APOD;SPTBN1;DSP;GABRP;<br>HMGCS1;DST;ACSL1;STAT1;LAMB2;SPINK5;FN1;PRRC2C;<br>RPSA;HSPG2;ENAH;TANC1;SFPQ;VCAN;COL3A1;PTPRC;<br>DES;NCL;CLIP4;COL5A2;SPARCL1;ITGA6;FGFR3;PTMA;SPRR1B;IVL |
| 19 | Vitinoin | 15 | FLG;TNXB;HMGCS1;FN1;DHCR24;HMGCR;VCAN;PTPRC;<br>SOAT1;COL5A2;PKP1;COL6A3;FGFR3;SPRR1B |
| 37 | Quercetin | 30 | EPAS1;UBR4;ITPR3;MTR;ENO1;ATP1A1;HMGCR;CLDN1;MYLK;LMNA;<br>CAPN2;ANXA6;TP63;SPTBN1;TNS1;MAP4K4;CD74;HMGCS1;STAT1;<br>FN1;RPSA;DHCR24;NEB;ENAH;COL3A1;SOAT1;COL7A1;TXNIP;ITGA6;FGFR3 |

In Figure 7, two subgraphs obtained from PPI networks are shown, including the following 25 genes: COL6A3, COL3A1, COL5A2, COL7A1, ITGA6, FN1, AGRN, HSPG2, VCAN, TTN, SOAT1, HSP90AB1, ATP6V1B1, HMGCR, HMGCS1, DSP, HRNR, FLG2, FLG, PKP1, RPL35A, RPS27A, RPS14, RPSA, and RPL37A. Some of these genes have been reported to be related to skin diseases. For example, genes related to the extracellular matrix shown in Figure 8 (including COL6A3, COL3A1, COL5A2, COL7A1, ITGA6, FN1, AGRN, HSPG2, VCAN, DSP) directly contribute to skin cutaneous diseases, such as LP [54–56]. Mutations in genes related to ribosomes (RPL35A, RPS27A, RPS14, RPSA, and RPL37A) are the key to several disorders [57–61]. These results may play a crucial role in identifying putative therapeutic targets and thereby identifying effective drugs.

**Figure 7:**
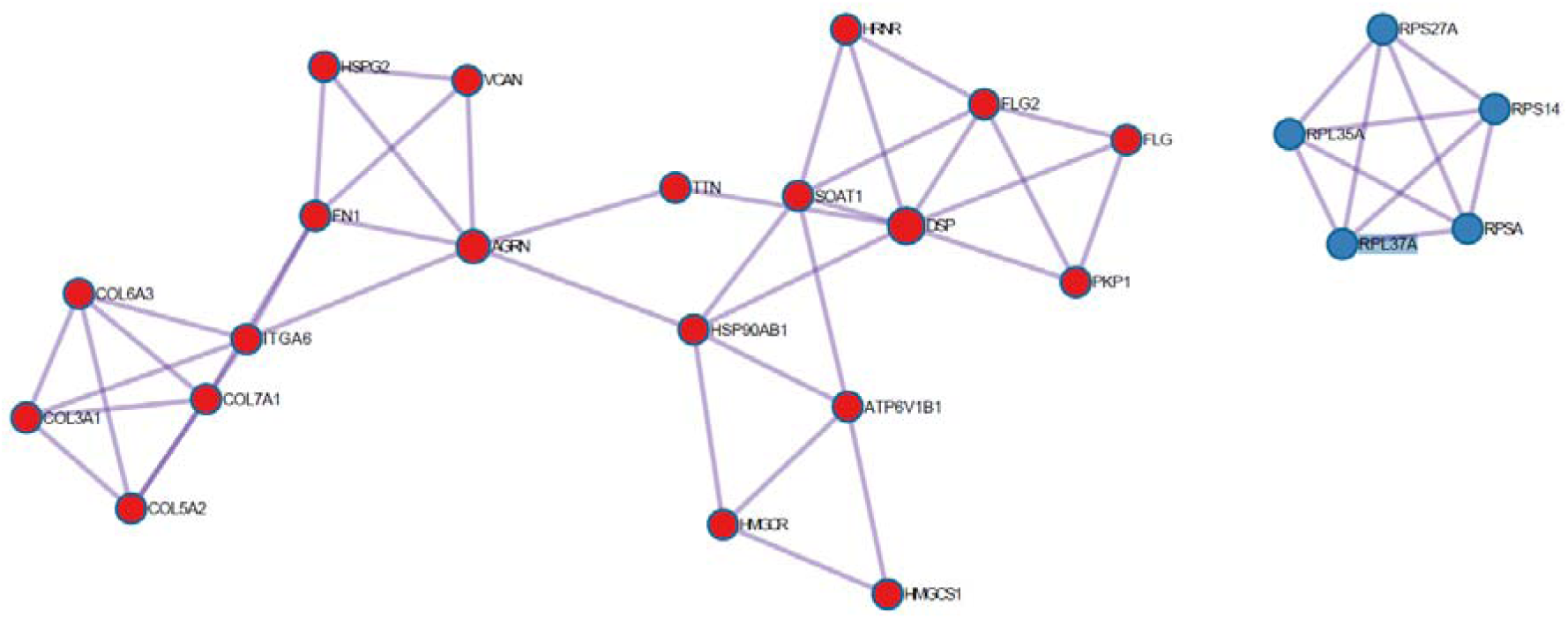
Two highly interconnected clusters in a protein-protein interaction based on 100 genes of ssvm3 provided to Metascape enrichment analysis.

**Figure 8:**
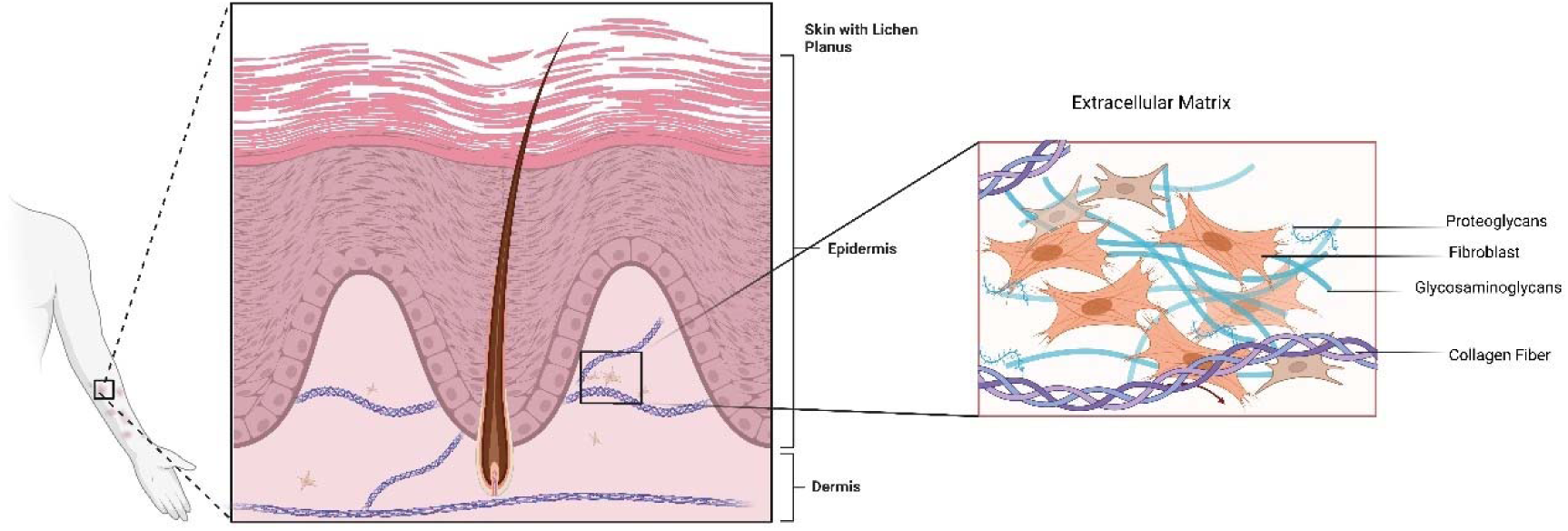
Skin of a patient with lichen planus affecting the arm area. Critical expressed genes in Figure 7 are related to skin dermis presented here. Figure created with BioRender.com.

In Figure 9, four transcription factors were reported (i.e., RELA, STAT3, TP53, STAT1). TP53 has been reported to be overexpressed in OLP [16, 62] and related to malignant transformation in OLP [63]. RELA (also called p65 or NF-κB/p65) has been reported to be overexpressed in LP, suggesting a role in disease development [64–66]. Activation of STAT1 has been observed in the skin epidermis of LP patients [19]. STAT3 mutations are related to autoimmune diseases including skin diseases [67]. These identified transcription factors can act as therapeutic targets in LP.

**Figure 9:**
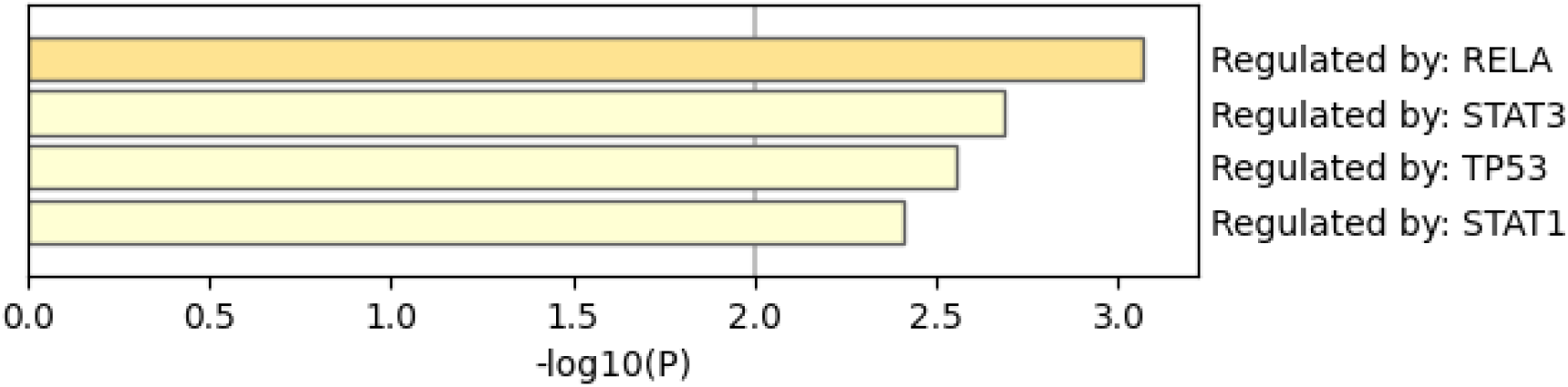
Four transcription factors provided by Metascape according to the list of 100 input genes of ssvm3.

#### 3.2.3 Dataset3

We uploaded genes obtained from each method to Enrichr. Table 8 shows disease-gene association, while Table 9 presents tissue terms according to expressed genes. From Table 8, we can see that ssvm1 and ssvm2, followed by ssvm3, generated the best results according to 3 out of 38 genes expressed in lichen planus. For autoimmune diseases and dermatologic diseases terms, ssvm3 performed the best, as 15 genes were expressed in autoimmune diseases and 10 genes were expressed in dermatologic disorders. Although ssvm1, ssvm2, and ssvm3 ranked 34th for lichen planus, they had the same number of expressed genes and were far better in terms of the ranking and number of expressed genes when considering the other two terms. In Table 9, our methods demonstrated superior results when compared to the baseline methods. Specifically, the skin tissue term was ranked 1st by ssvm1 and ssvm2, while it was ranked 3rd by ssvm3. Moreover, the number of expressed genes was 52 out of 2316 for ssvm1 and ssvm2, outperforming ssvm3. However, results were significant for all our methods. SAM ranked the skin tissue as the 1st term, but the number of expressed genes was 25 less than those obtained via our methods. Figure 10 provides a visualization of gene intersection among all methods. It should be noted that ssvm3 had 52 genes differing from all other methods and that ssvm1 and ssvm2 shared 37 genes. Together, all our methods shared 28 genes. SAM shared 13 genes with our methods. These results demonstrate that there was not a total overlapping among all methods, as each method produced a different list of genes affecting the enrichment analysis results. In Supplementary Data Sheet 3, we provide each list of genes produced via each method used in enrichment analysis. Moreover, we provide enrichment analysis results pertaining to Tables 8 and 9 in Supplementary Table 3_A and Table 3_B, respectively.

**Figure 10:**
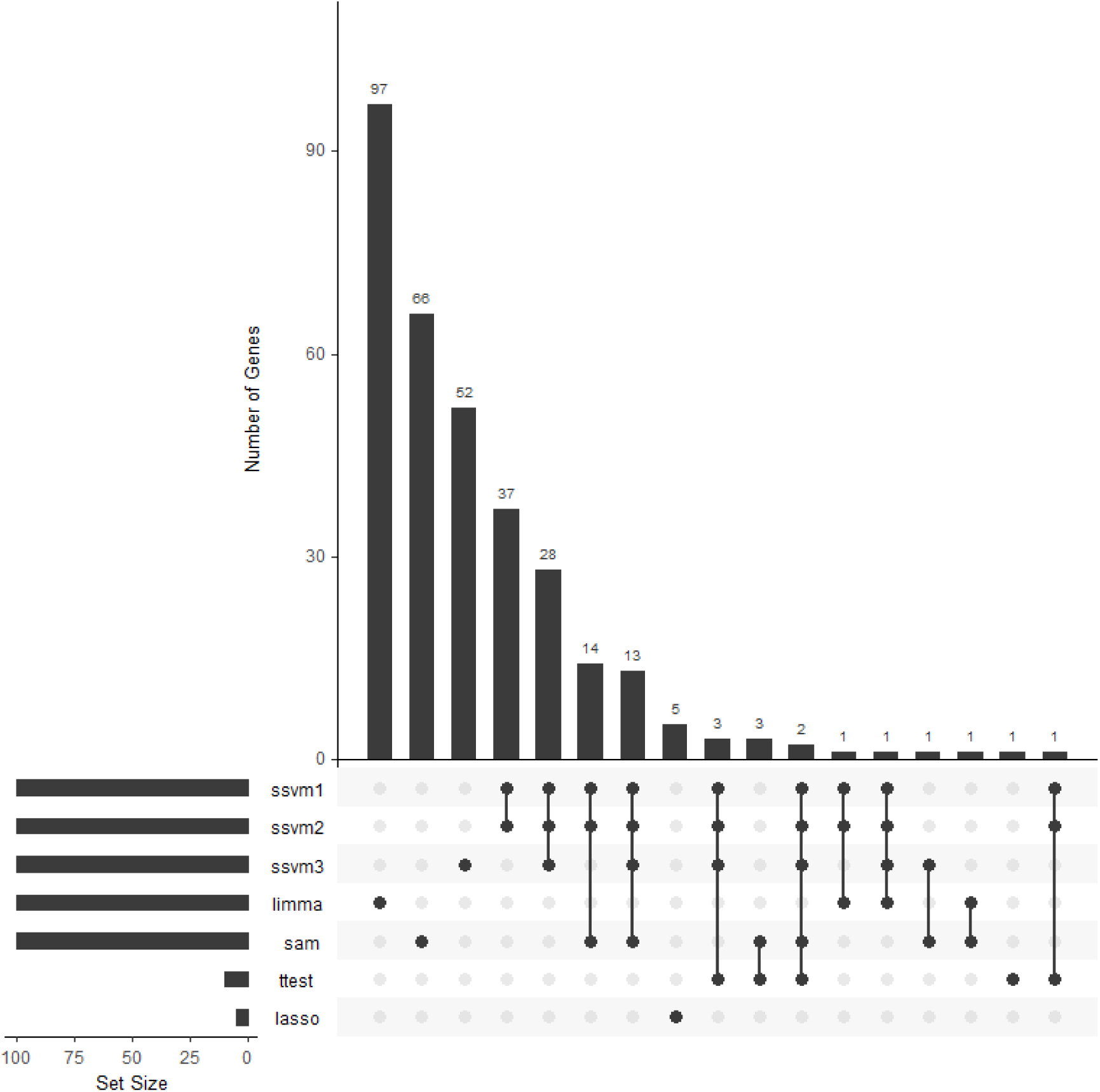
Visualization of intersecting sets pertaining to each gene list provided by each method when Dataset3 was used.

**Table 8:** Retrieved enriched terms within “DisGeNET” via Enrichr according to uploaded genes via methods using Dataset3, showing association between human genes and diseases. Rank column shows the order of terms when retrieved. Overlap displays a ratio-like format, corresponding to the number of uploaded genes overlapping to those in the terms. The best result of each method is in bold.

| Method | Rank | Term | Overlap | *P*-value | Adjusted *P*-value |
| --- | --- | --- | --- | --- | --- |
| ssvm1 | 34 | LICHEN PLANUS | **3/38** | $9.01 \times 10^{-4}$ | $6.40 \times 10^{-2}$ |
| | 2121 | AUTOIMMUNE DISEASES | 6/1060 | $4.38 \times 10^{-1}$ | $4.99 \times 10^{-1}$ |
| | 37 | DERMATOLOGIC DISORDERS | 8/406 | $9.84 \times 10^{-4}$ | $6.40 \times 10^{-2}$ |
| ssvm2 | 34 | LICHEN PLANUS | **3/38** | $9.01 \times 10^{-4}$ | $6.40 \times 10^{-2}$ |
| | 2121 | AUTOIMMUNE DISEASES | 6/1060 | $4.38 \times 10^{-1}$ | $4.99 \times 10^{-1}$ |
| | 37 | DERMATOLOGIC DISORDERS | 8/406 | $9.84 \times 10^{-4}$ | $6.40 \times 10^{-2}$ |
| ssvm3 | 59 | LICHEN PLANUS | 3/38 | $9.01 \times 10^{-4}$ | $3.80 \times 10^{-2}$ |
| | 33 | AUTOIMMUNE DISEASES | **15/1060** | $2.43 \times 10^{-4}$ | $1.81 \times 10^{-2}$ |
| | 11 | DERMATOLOGIC DISORDERS | **10/406** | $3.62 \times 10^{-5}$ | $7.52 \times 10^{-3}$ |
| limma | - | LICHEN PLANUS | - | - | - |
| | 1160 | AUTOIMMUNE DISEASES | 5/1060 | $6.16 \times 10^{-1}$ | $7.34 \times 10^{-1}$ |
| | 1327 | DERMATOLOGIC DISORDERS | 1/406 | $8.72 \times 10^{-1}$ | $9.08 \times 10^{-1}$ |
| sam | - | LICHEN PLANUS | - | - | - |
| | 1142 | AUTOIMMUNE DISEASES | 8/1060 | $1.60 \times 10^{-1}$ | $3.20 \times 10^{-1}$ |
| | 1724 | DERMATOLOGIC DISORDERS | 3/406 | $3.31 \times 10^{-1}$ | $4.42 \times 10^{-1}$ |
| *t*-test | 24 | LICHEN PLANUS | 1/38 | $1.88 \times 10^{-2}$ | $8.83 \times 10^{-2}$ |
|  | - | AUTOIMMUNE DISEASES | - | - | - |
| | 21 | DERMATOLOGIC DISORDERS | 2/406 | $1.66 \times 10^{-2}$ | $8.83 \times 10^{-2}$ |
| lasso | - | LICHEN PLANUS | - | - | - |
| | 271 | AUTOIMMUNE DISEASES | 1/1060 | $2.38 \times 10^{-1}$ | $2.68 \times 10^{-1}$ |
|  | - | DERMATOLOGIC DISORDERS | - | - | - |

**Table 9:** Retrieved enriched terms within “ARCHS4 Tissues” via Enrichr according to uploaded genes via methods using Dataset3, showing genes expressed within tissue types. Rank column shows the order of terms when retrieved. Overlap displays a ratio-like format, corresponding to the number of uploaded genes overlapping to those in the terms. The best result of each method is in bold.

| Method | Rank | Term | Overlap | *P*-value | Adjusted *P*-value |
| --- | --- | --- | --- | --- | --- |
| ssvm1 | 1 | SKIN (BULK TISSUE) | 52/2316 | $3.97 \times 10^{-23}$ | $4.28 \times 10^{-21}$ |
| ssvm2 | 1 | SKIN (BULK TISSUE) | 52/2316 | $3.97 \times 10^{-23}$ | $4.28 \times 10^{-21}$ |
| ssvm3 | 3 | SKIN (BULK TISSUE) | 34/2316 | $2.95 \times 10^{-9}$ | $1.06 \times 10^{-7}$ |
| limma | 50 | SKIN (BULK TISSUE) | 10/2316 | $7.34 \times 10^{-1}$ | $9.99 \times 10^{-1}$ |
| sam | 1 | SKIN (BULK TISSUE) | 25/2316 | $1.40 \times 10^{-4}$ | $1.52 \times 10^{-2}$ |
| *t*-test | 8 | SKIN (BULK TISSUE) | 5/2316 | $3.16 \times 10^{-3}$ | $3.35 \times 10^{-2}$ |
| lasso | 54 | SKIN (BULK TISSUE) | 1/2316 | $4.59 \times 10^{-1}$ | $4.59 \times 10^{-1}$ |

We selected ssvm3 for further biological analysis, as it achieved good results. In Table 10, we retrieve drug-genes association. The three drugs (Retinoic acid, Vitinoin, and Alitretinoin) coincide with those in Tables 4 and 7, demonstrating improvements in treatment in clinical studies [36–39]. In Supplementary Table 3_C, we provide enrichment analysis results pertaining to drug terms within DSigDB

**Table 10:**
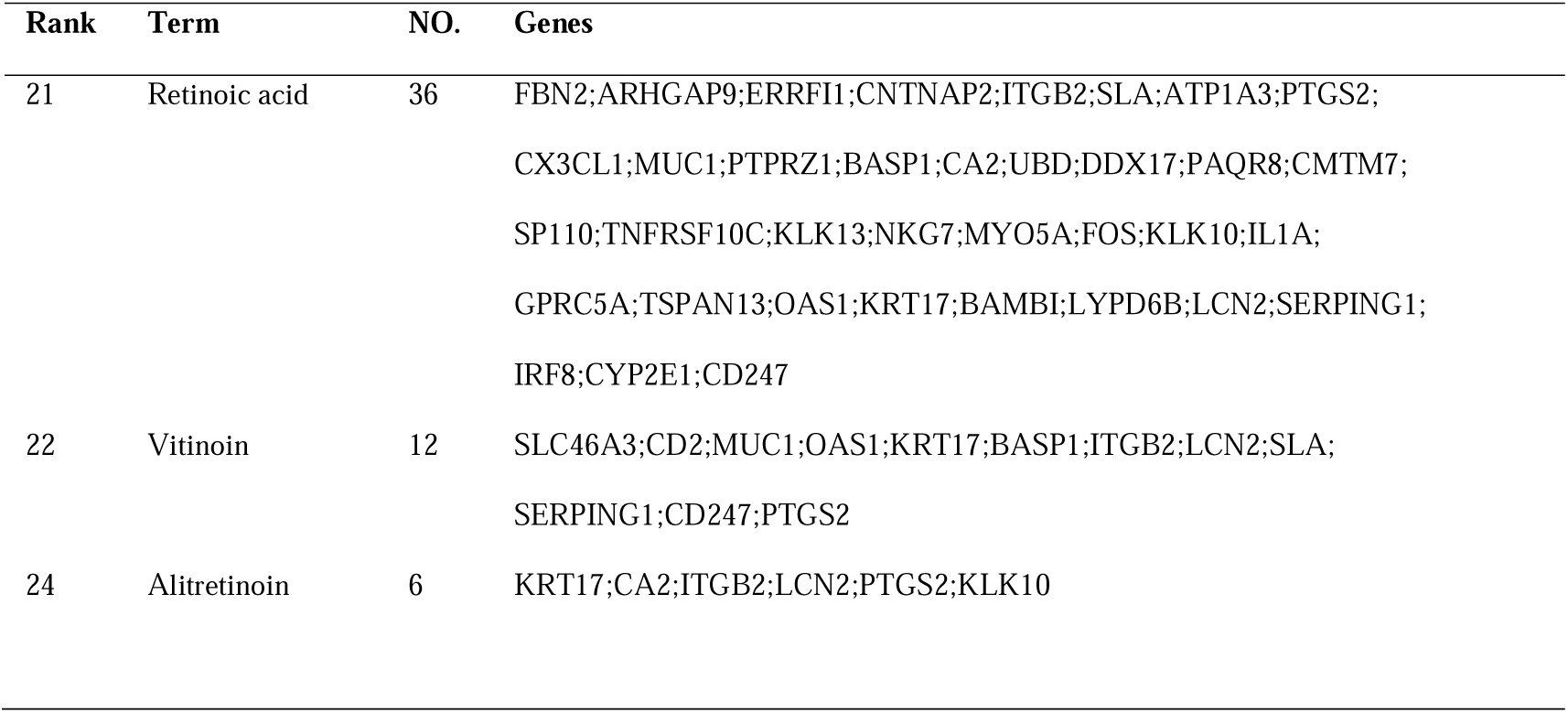
Retrieved enriched terms within “DSigDB” via Enrichr according to uploaded genes using Dataset3, showing genes (column: Genes) associated with drugs (column: Term). Rank column shows the order of terms when retrieved. NO. indicates the number of genes.

Regarding the two subgraphs in Figure 11 obtained from the PPI network, the three genes (PI3, LCE2A, LCE2D) are related to the formation of the cornified envelope within the startum corneum of the skin epidermis (Figure 12), while the LCP1 and ITGB2 are related to cytokine signalling in immune system, indicating their inflammatory role in the formation and maintenance of skin cornified envelope in LP [68–70].

**Figure 11:**
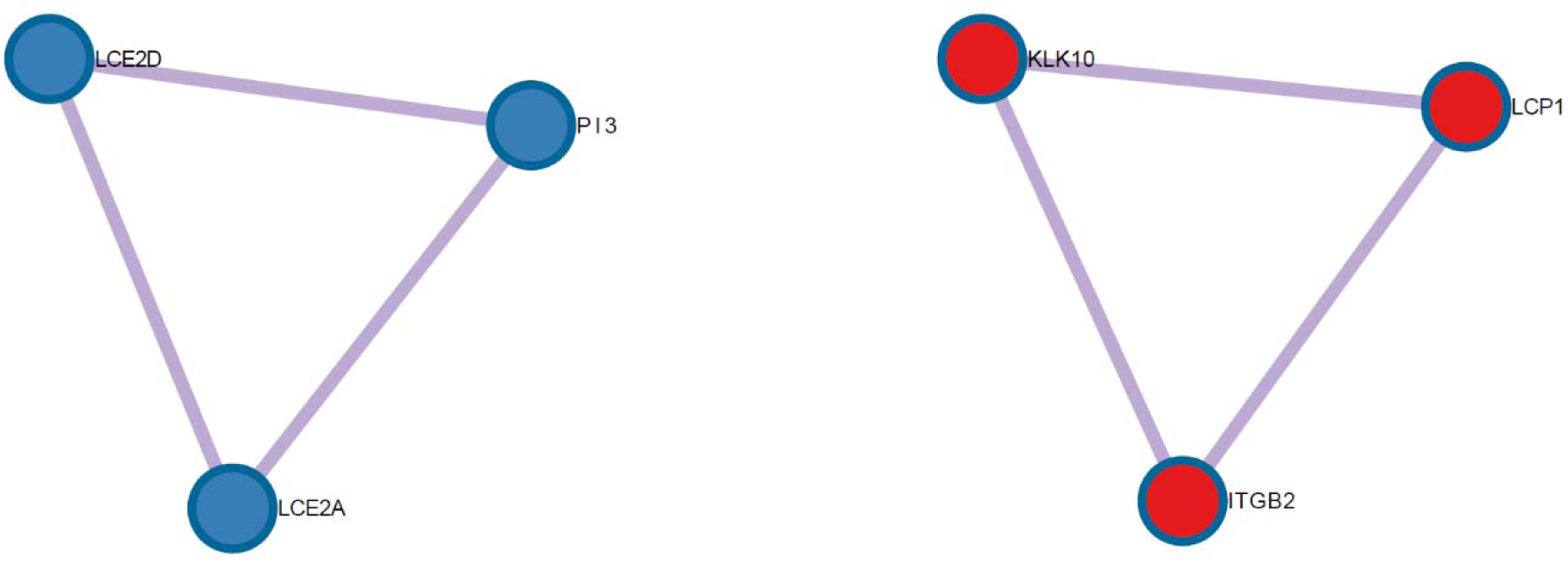
Two highly interconnected clusters in a protein-protein interaction based on 100 genes of ssvm3 provided to Metascape enrichment analysis.

**Figure 12:**
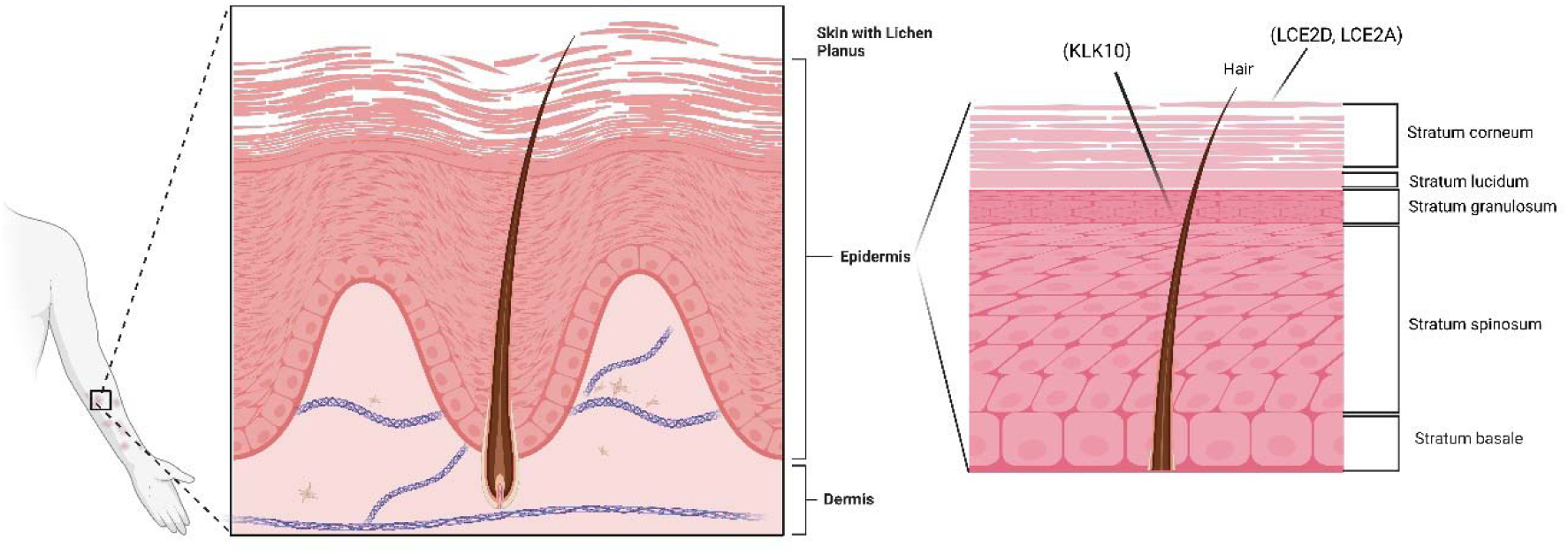
Skin of a patient with lichen planus affecting the arm area. Critical expressed genes obtained from Figure 11 are related to the skin epidermis layers. Figure created with BioRender.com.

Figure 13 lists 11 transcription factors, STAT1, STAT3, STAT6, RELA, USF2, NFKB1, FOS, USF1, HDAC1, CREB1, and AR. As in Dataset2, STAT1, STAT3, and RELA have been reported to be associated with LP [19, 64–67]. HDAC1 plays a key role in skin inflammation through Filaggrin expression, which is part of skin barrier dysfunction [71]. These transcription factors can be useful therapeutic targets in inflammatory skin diseases, such as LP.

**Figure 13:**
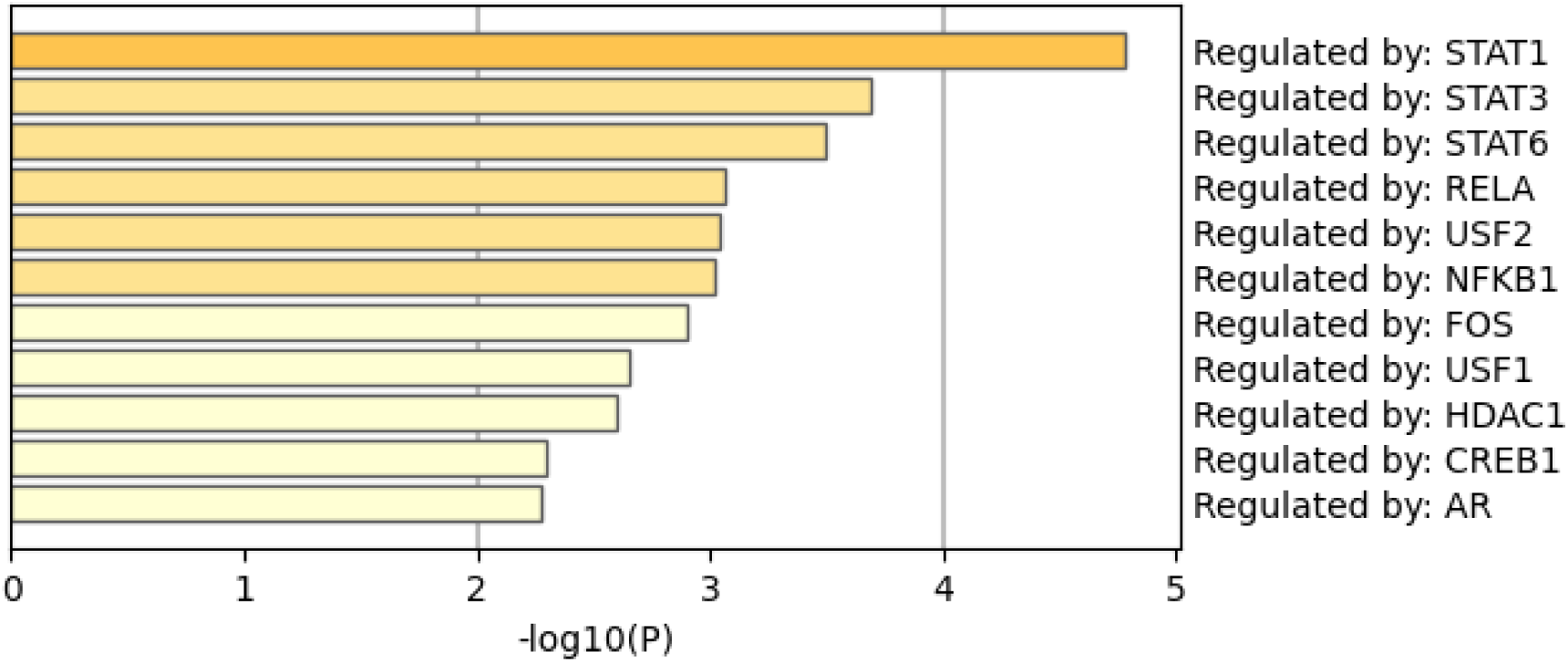
Eleven transcription factors provided by Metascape according to the list of 100 input genes of ssvm3.

#### 3.2.4 Sparse Models Explanation

##### Dataset1

In Figure 14, we provide insights pertaining to models used in the Results section. Figures 14(a-d) show that there were statistically significant differences in predictions of healthy controls and LP patients (*p-*value of all models << 0.01, obtained from a t-test), indicating that these models applied to Dataset1 are expected to be general predictors to healthy controls and LP. In Figure 14(e), ssvm1 and ssvm2 achieved an AUC of 0.983, while ssvm3 and lasso both achieved an AUC of 1.00.

**Figure 14:**
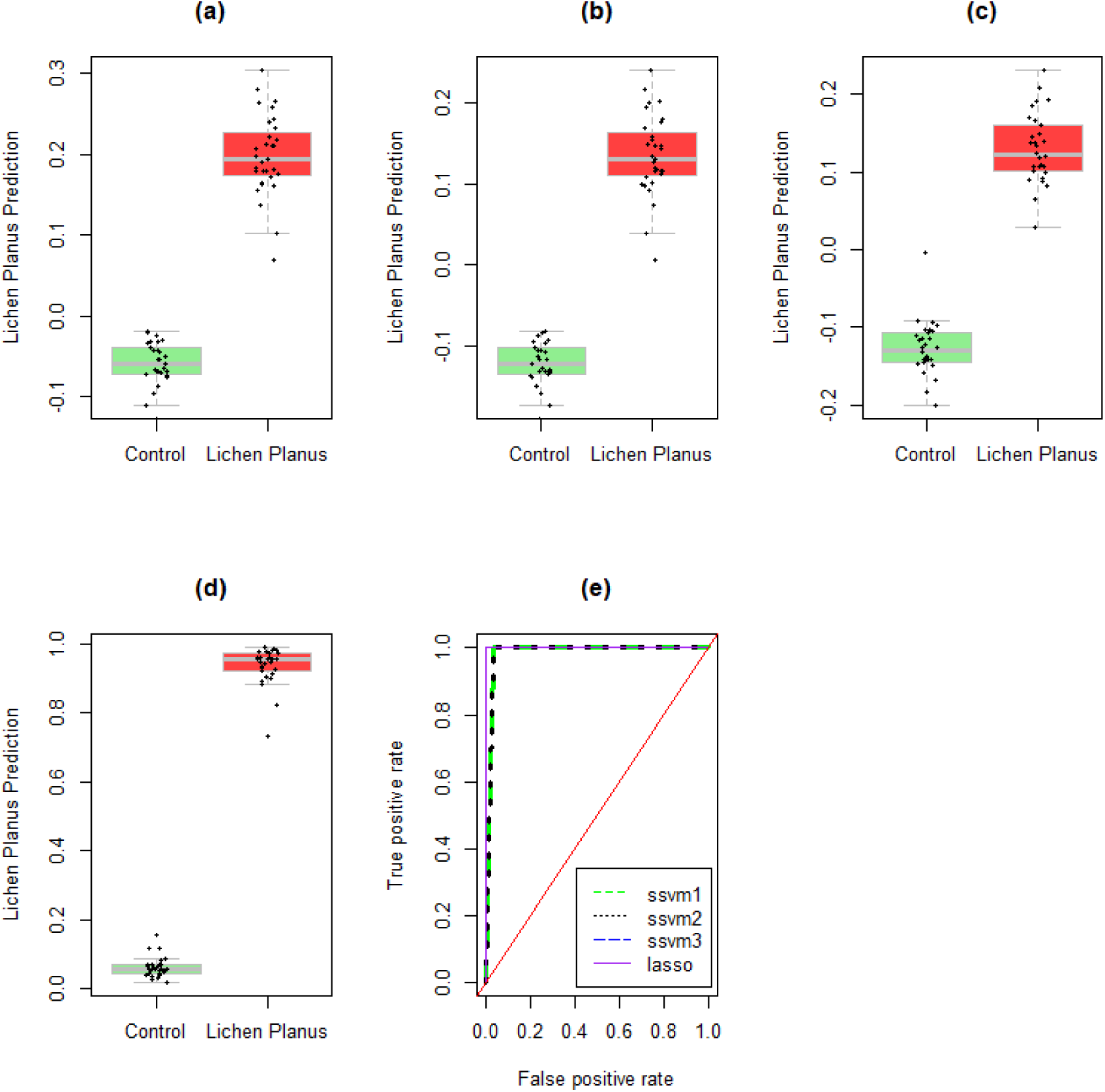
Prediction results regarding healthy controls and lichen planus (LP) when Dataset1 was used. Boxplot and strip charts (a-d) show the differences in predicted LP pertaining to healthy individuals and individuals with LP for ssvm1, ssvm2, ssvm3, and lasso, respectively. (e) demonstrates the ROC curves by displaying the true positive rate along the y-axis and the false positive rate along the x-axis.

Figure 15 shows that our methods (Figures 15(a-c)) reduced more weights of genes to zero compared to lasso in Figure 15(d). These results demonstrate that our optimized methods led to sparser feature vector representations than lasso.

**Figure 15:**
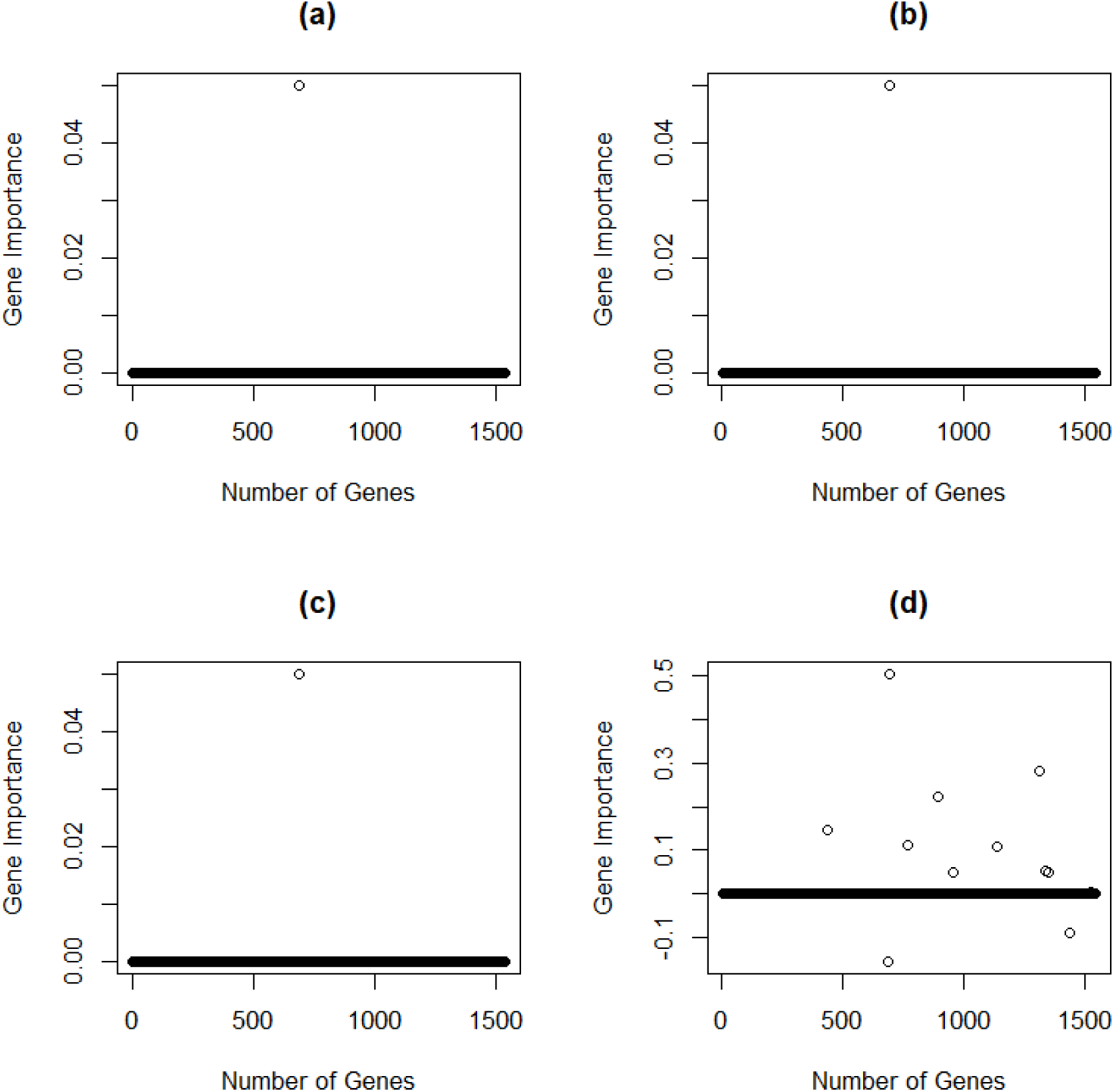
Gene importance of all genes pertaining to Dataset1 produced via ssvm1 (a), ssvm2 (b), ssvm3 (c), and lasso (d).

##### Dataset2

In Figure 16, we provide an explanation for models utilized in the Results section. Figures 16(a-d) show that differences in predictions of healthy controls and LP patients were statistically significant (*p-*value of all models << 0.01, obtained from a t-test), indicating that these models applied to Dataset2 are not expected to be disease specific. In Figure 16(e), all models achieved an AUC of 1.00. Figure 17 shows that our methods (Figures 17(a-c)) shrunk weights of genes to zero, while lasso (Figure 15(d)) shrunk more coefficients of genes to zero than our methods. These results demonstrate sparsity induced via each model.

**Figure 16:**
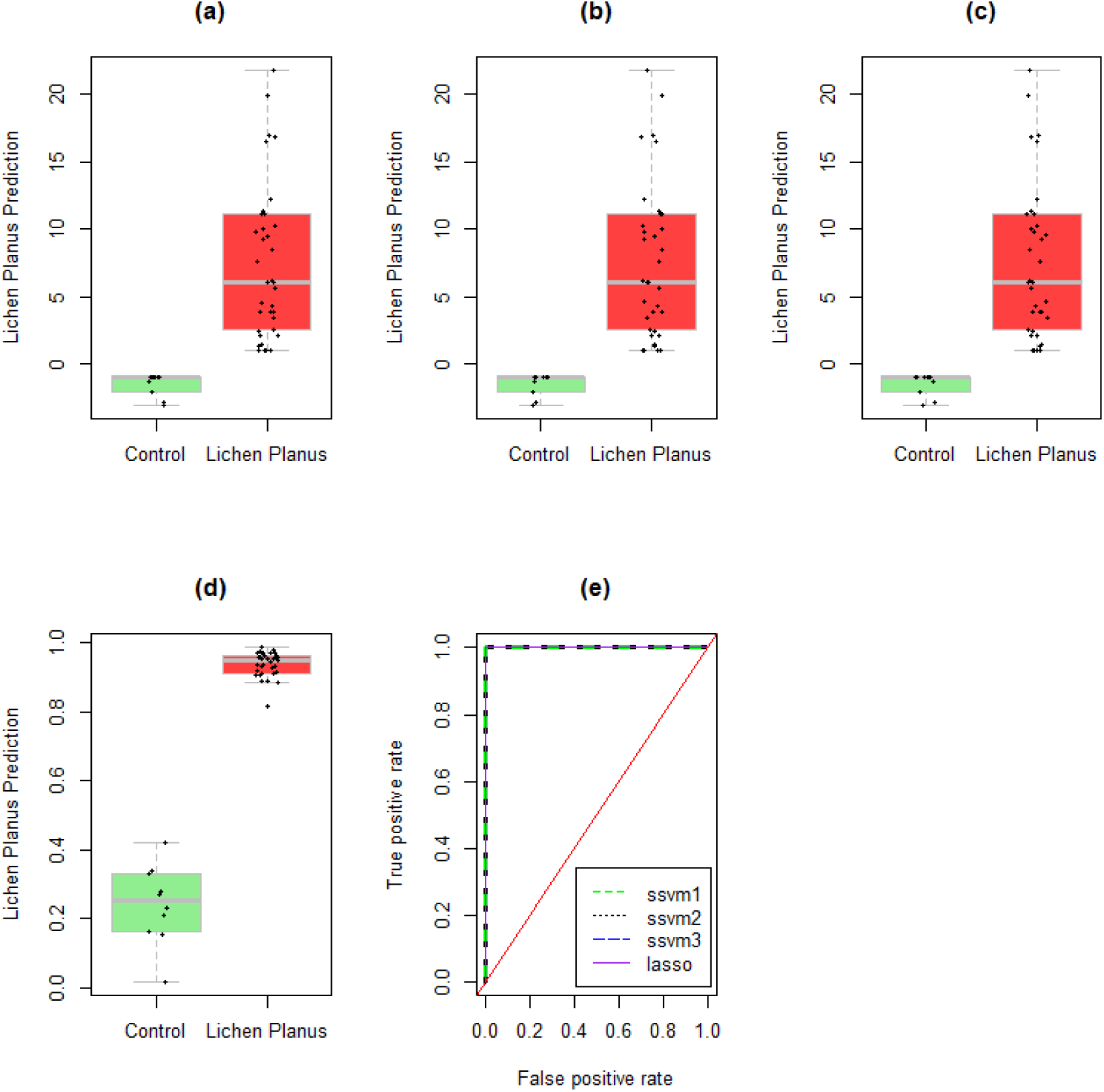
Prediction results regarding healthy controls and lichen planus when Dataset1 was used. Boxplot and strip charts (a-d) show the differences in predicted lichen planus pertaining to healthy individuals and individuals with LP for ssvm1, ssvm2, ssvm3, and lasso, respectively. (e) Demonstrates the ROC curves by displaying the true positive rate along the y-axis and the false positive rate along the x-axis.

**Figure 17:**
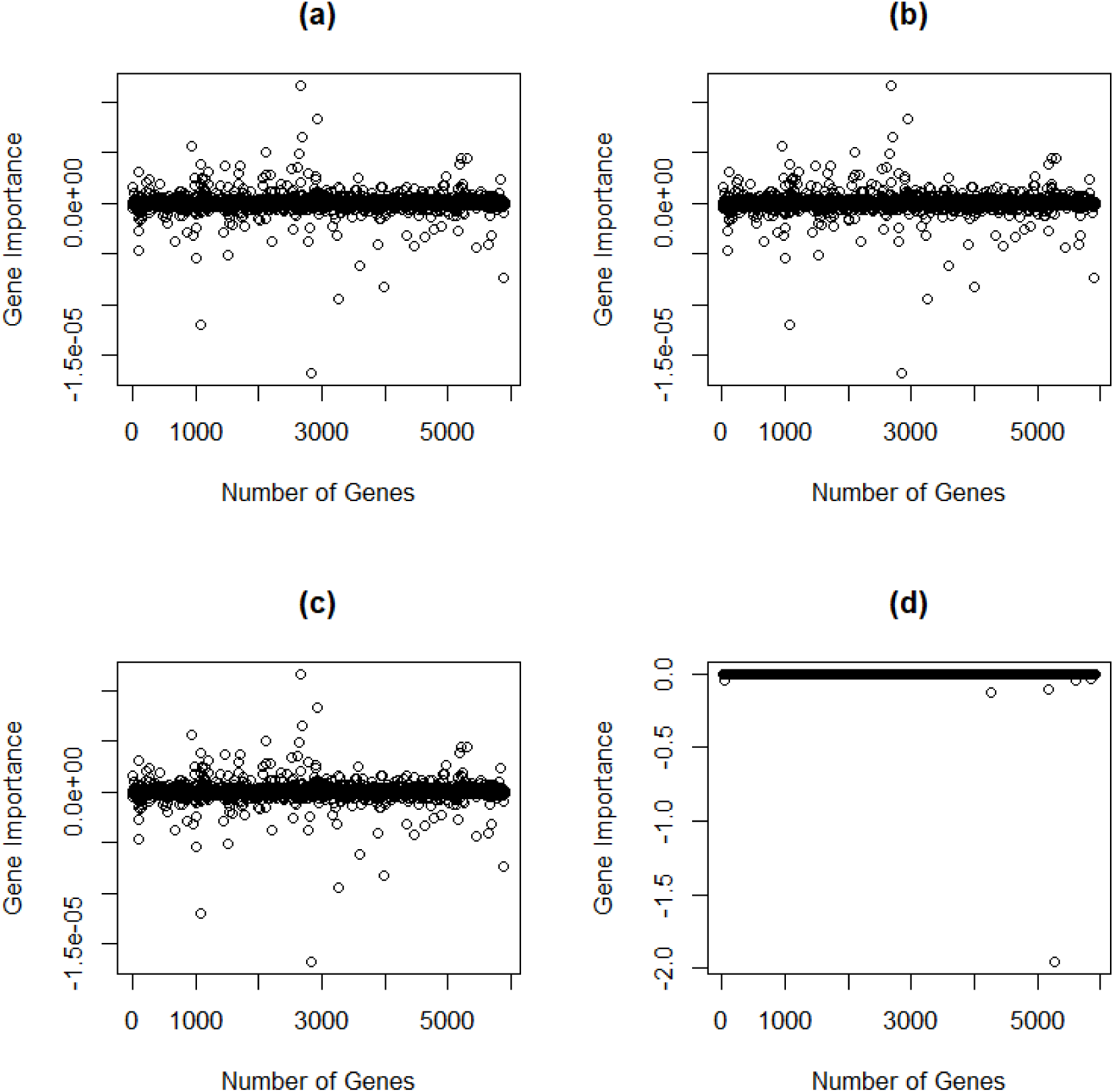
Gene importance of all genes pertaining to Dataset2 produced via ssvm1 (a), ssvm2 (b), ssvm3 (c), and lasso (d).

##### Dataset3

In Figure 18, we provide interpretation for models employed in the Results section. Figures 18(a-d) show that differences in predictions of healthy controls and LP patients were statistically significant (*p-*value of all models << 0.01, obtained from a t-test), showing that these models applied to Dataset3 are expected to be general predictors of LP and healthy controls. In Figure 18(e), our models achieved an AUC of 0.928, while lasso had an AUC of 1.00.

**Figure 18:**
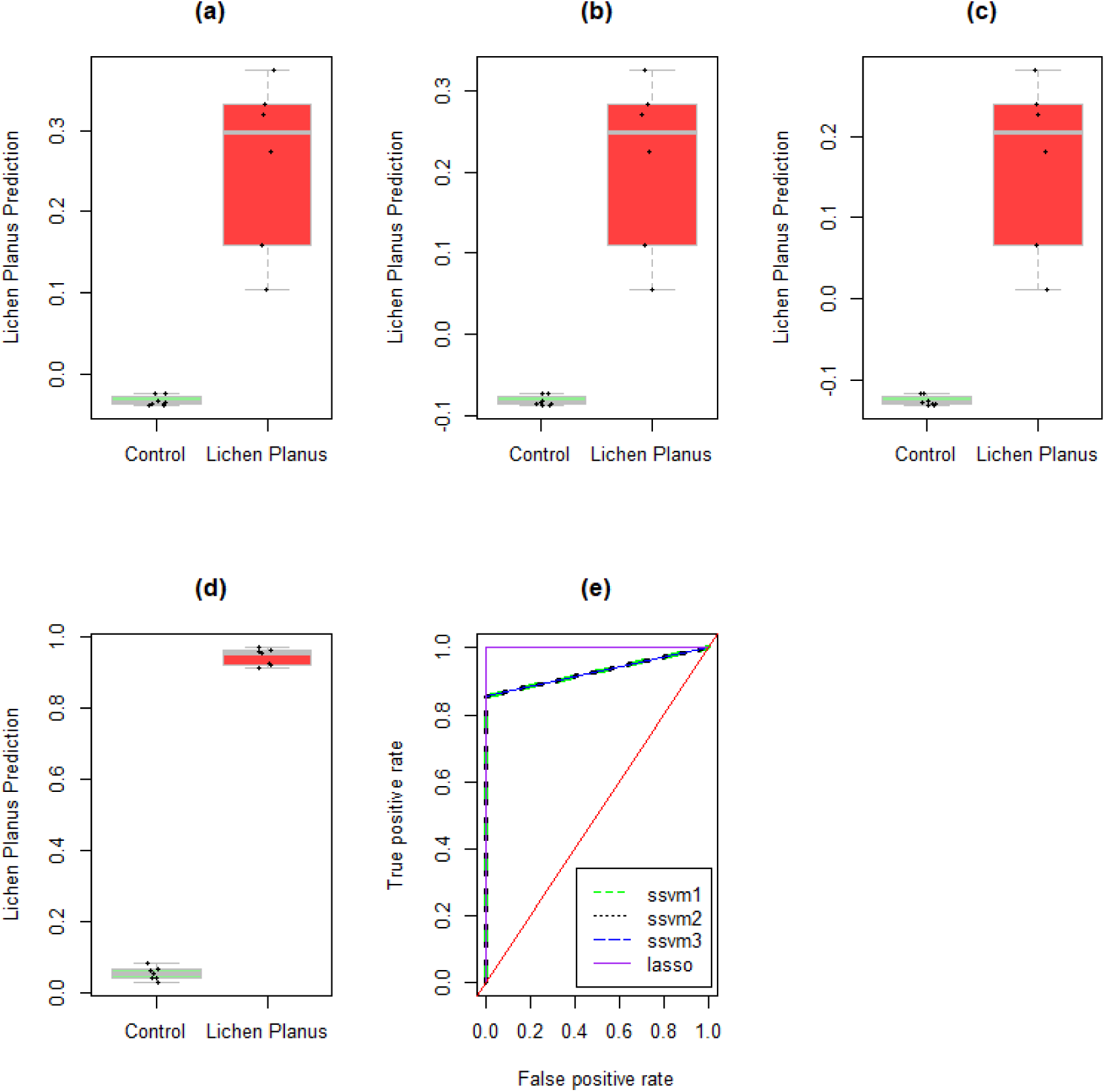
Prediction results regarding healthy controls and lichen planus when Dataset3 was used. Boxplot and strip charts (a-d) show the differences in predicted LP pertaining to healthy individuals or individuals with LP for ssvm1, ssvm2, ssvm3, and lasso, respectively. (e) Demonstrates the ROC curves for each model by displaying the true positive rate along the y-axis and the false positive rate along the x-axis

As seen in Figure 19, our methods (Figures 19(a-c)) shrunk more weights of genes to zero compared to lasso (Figure 19(d)). These results demonstrate that our constrained optimization problem led to a sparser representation than that of lasso.

**Figure 19:**
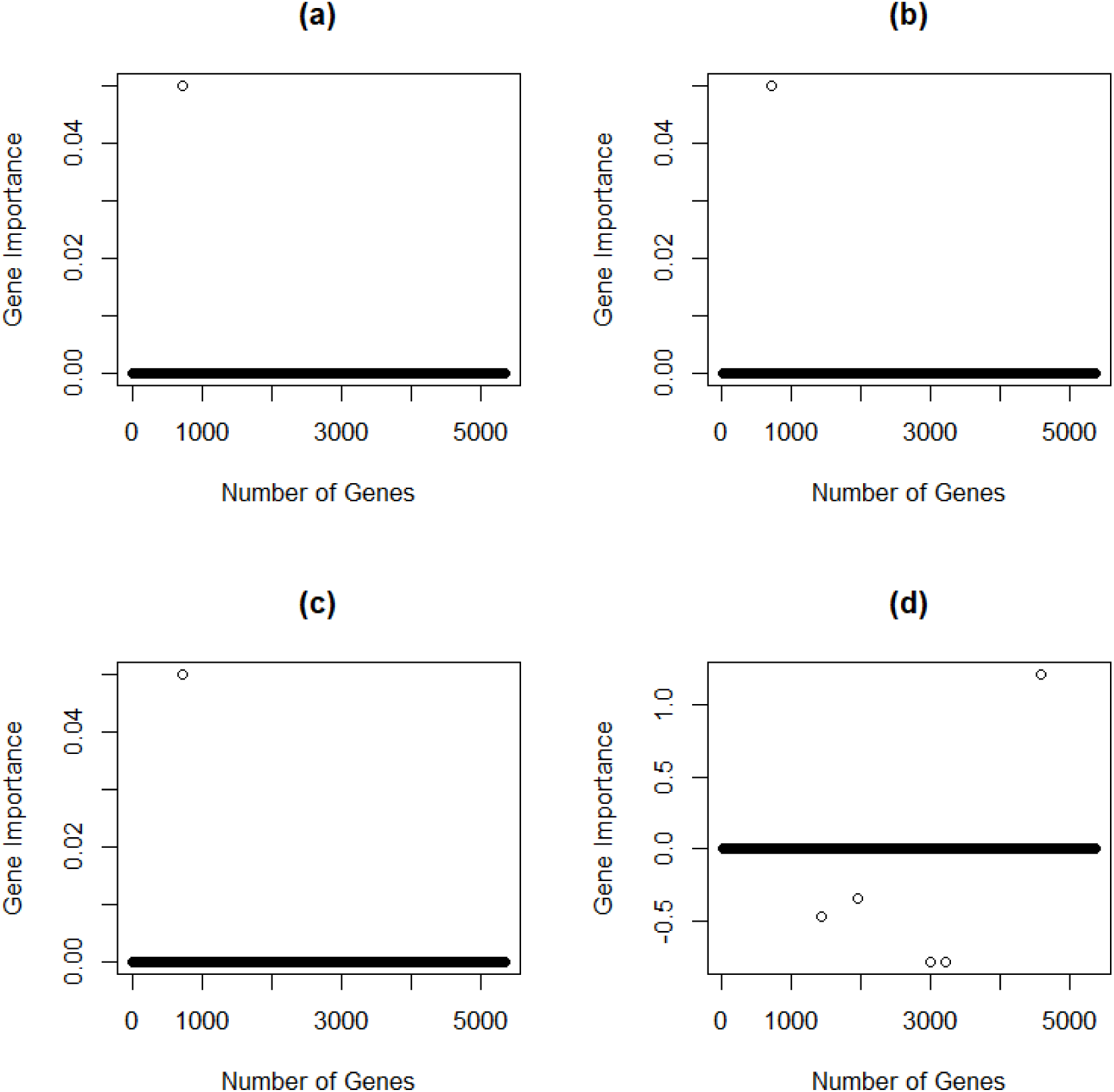
Gene importance of all genes pertaining to Dataset3, as produced via ssvm1 (a), ssvm2 (b), ssvm3 (c), and lasso (d).

## 4 Discussion

Identifying influential genes and relevant drugs contributes to a greater understanding of the molecular mechanism of LP as well as of disease treatment. As a result, we presented a computational framework working as follows: first, we introduced three sparse optimization problems pertaining to SVM, named ssvm1, ssvm2, and ssvm3. We provide each gene expression dataset as input to our methods, solved using CVXR solver in R. The output of each method was a weight vector w, corresponding to the importance of each gene in the provided dataset. Genes associated with higher weights indicated the importance of these genes. We ordered and then selected the top 100 genes according to the top 100 weights in w. Then, we provided output genes via each method to Enrichr and Metascape to perform enrichment analysis. With Enrichr, our methods (ssvm1, ssvm2, and ssvm3) achieved the best results as compared to baseline methods, and the retrieved terms had a greater number of expressed genes in disease terms, which included lichen planus, autoimmune diseases, dermatologic disorders, and skin diseases. Regarding tissue types, our methods also outperformed all baseline methods, having more expressed genes within the tissue type term (i.e., skin tissue). Furthermore, we identified drugs associated with genes via Enrichr. With Metascape, we identified important genes as well as TFs related to the skin, which have been reported to play a key role in the pathogenesis and treatment of lichen planus.

As shown in the presented results (See Tables 2-4, 5-7, and 8-10), our computational framework consists of two parts: machine learning and enrichment analysis. Specifically, machine learning identified important genes within gene expression datasets, which were then provided to enrichment analysis tools to retrieve terms related to drugs, diseases, tissues, PPI networks, and TFs related to LP. In a clinical trial setting, these obtained results could help doctors to identify drugs, expressed genes, and therapeutic targets in T-cell mediated disease, such as lichen planus, which is incurable and for which current treatments are aimed at disease management due to the unknown etiology.

As clinical trials involve testing already approved drugs or new drugs, our computational framework enables the narrowing of the search for such drugs by providing a subset of effective drugs [72]; it also can narrow the search for putative therapeutic targets, thereby speeding up the drug discovery. It is worth noting that drugs associated with the genes in Tables 4, 7, and 10 could help clinicians personalize drug treatment. In other words, when these genes (or drug targets) are expressed, the corresponding drug could be a more suitable treatment choice [73]. This demonstrates the benefit of AI tools in clinical trials.

In the “Sparse Models Explanation” section, we demonstrate important information pertaining to studied sparse models, including ssvm1, ssvm2, ssvm3, and lasso. Specifically, Figures 14, 16, and 18 demonstrate the statistical significance of induced models, which are expected to acquire a good generalization when discriminating between the two classes of healthy controls and LP. However, Figures 15, 17, and 19 explored induced sparsity, in which genes of weak discriminative power were set to zero. It can be observed that the additional constraint IIW II_1_ ≤ λ enforced sparsity. Our models (i.e., ssvm1, ssvm2, and ssvm3) are based on SVM and therefore rely on the margin theory, in which the aim is to maximize the margin 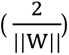, which is equivalent to minimizing the margin 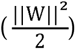 subject to additional constraints, as shown in Equation 1-4. The weight vector w provides importance of features and is derived based on the margin notion. However, lasso depends on linear models, including L1 penalty, to shrink coefficients to zero. According to our results in Section 3.2, results derived from our models relying on margin theory are better than those depending on linear models, such as lasso [74–76].

We set parameters for the four methods as follows: ssvm1(*c* = 2, *k*=1, λ=0.05), ssvm2(*c* = 2, *k*=1, λ=0.05, *r* = 0.1), ssvm3(*c* = 2, *k*=2, λ=0.05, *r* = 0.1), lasso(λ=0.05). As the output of each of our methods was 100, the probability for other results coinciding with ours in the three datasets are 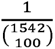, 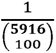, and 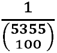, respectively; 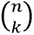 denotes the binomial coefficient, where *n* is the number of genes in the studied dataset and *k* refers to the number of genes selected by our method. Therefore, it is very unlikely that the results of other methods coincide with ours.

## 5 Conclusions and Future Work

Here, we introduce a computational framework consisting of machine learning coupled with enrichment analysis to identify important genes and drugs related to LP. The input to our framework was a gene expression dataset pertaining to healthy controls and LP patients; we downloaded three gene expression datasets from the gene expression omnibus (GSE63741, GSE193351, GSE52130). In each processed dataset, each sample was associated with a binary label (i.e., healthy control or LP). We then provided the input dataset to three machine learning-based methods. All methods were new, sparse versions of SVM. The output of each method was a weight vector w. We selected the top *p* genes based on the top *p* corresponding weights in w and input genes selected by our methods and other baseline methods to enrichment analysis tools Enrichr and Metascape. Results show that our methods performed better than baseline methods in disease identification, showing more expressed genes and better identifying tissue type (i.e., skin tissue) with more expressed genes. With further enrichment analysis, we identified 5 drugs (e.g., dexamethasone, retinoic acid, and quercetin), 23 unique TFs (e.g., NFKB1, p65, HDAC1, STAT1, STAT3), and 45 unique genes (e.g., PSMB8, PSMB10, PSMB9, KRT31, KRT16, KRT19, KRT17, KRT15, COL6A3, COL3A1, LCE2D, LCE2A) that have been reported to be related to T cell-mediated autoimmune inflammatory mechanisms, therapeutic targets, and treatment of LP. Our methods are publicly available on the GENEvaRX web server at https://aibio.shinyapps.io/GENEvaRX/.

Future work includes: (1) adapting our computational framework to identify important expressed genes, therapeutics targets, and drugs in different skin diseases, such as psoriasis, atopic dermatitis, and lupus; (2) collaborating with hospitals to utilize our tool in clinical trials; and (3) identifying gene signatures from gene expression data using our methods.

**Turki Turki** received a B.S. in computer science from King Abdulaziz University, an M.S. in computer science from NYU.POLY, and a Ph.D. in computer science from the New Jersey Institute of Technology. He is currently an associate professor with the Department of Computer Science, King Abdulaziz University, Saudi Arabia. His research interests include machine learning, data science and bioinformatics. His research has been published in journals such as *Scientific Reports, IEEE Journal on Selected Topics in Signal Processing*, *Frontiers in Genetics and Computers in Biology and Medicine*. He is an editorial board member of *Sustainable Computing: Informatics and Systems, BMC Medical Genomics, PLOS ONE,* and *Computers in Biology and Medicine*.

**Y-h. Taguchi** received a B.S. degree in physics from the Tokyo Institute of Technology and a Ph.D. degree in physics from the Tokyo Institute of Technology. He is currently a full professor with the Department of Physics, Chuo University, Japan. His works have been published in leading journals such as *Physical Review Letters*, *Bioinformatics*, and *Scientific Reports*. His research interests include bioinformatics, machine learning, and nonlinear physics. He is also an editorial board member of *PloS ONE*, *BMC Medical Genomics*, *Medicine* (Lippincott Williams & Wilkins journal), *BMC Research Notes*, and *IPSJ Transaction on Bioinformatics*.

