## Supplementary material for "GENEvaRX: A Novel AI-Driven Method and Web Tool Can Identify Critical Genes and Effective Drugs for Lichen Planus": GENEvaRX_Screenshots.docx

- Go to <https://aibio.shinyapps.io/GENEvaRX/>

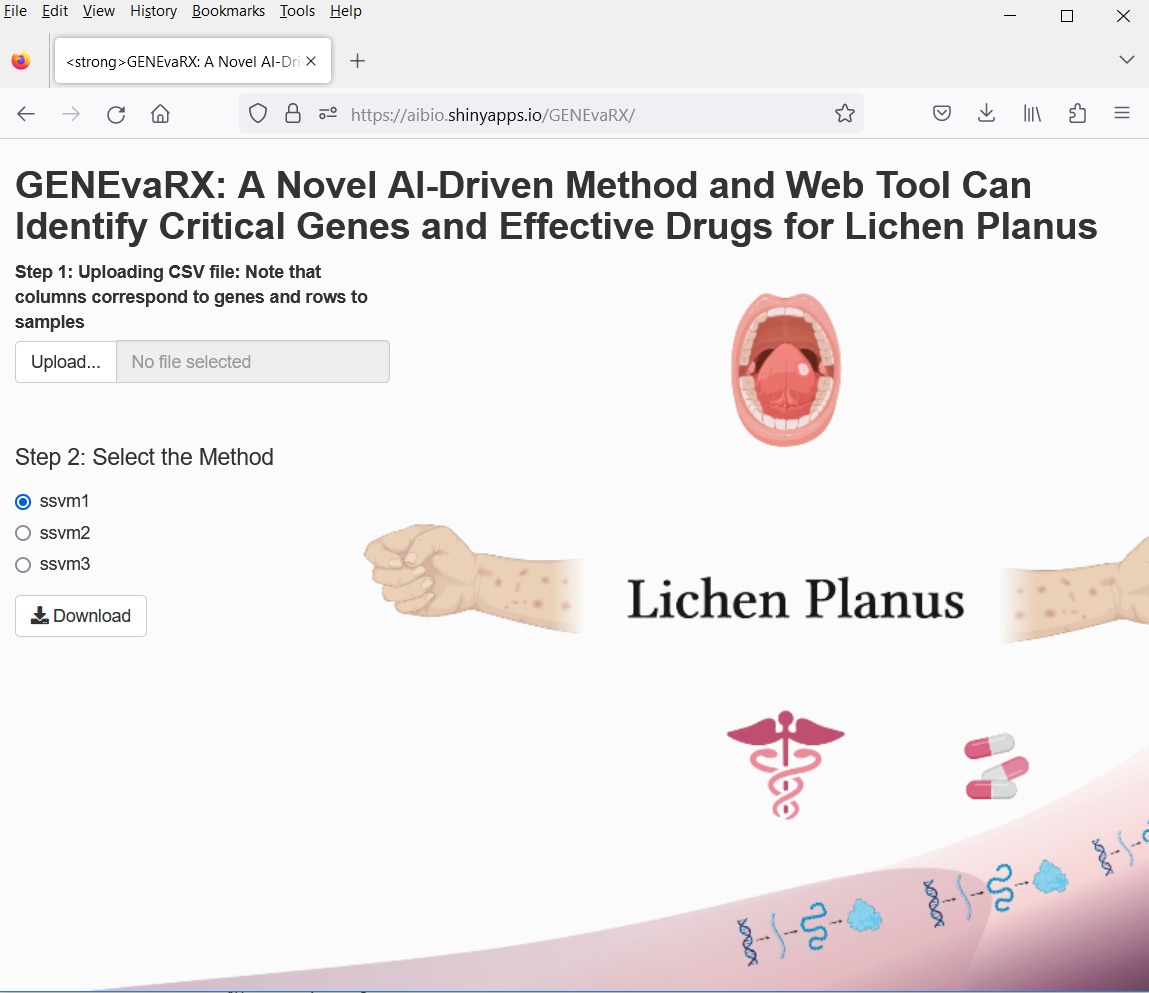

- Upload the dataset from Datasets folder within Supplementary Materials. Suppose we upload “GSE63741_Dataset1”. Then, we have the following.

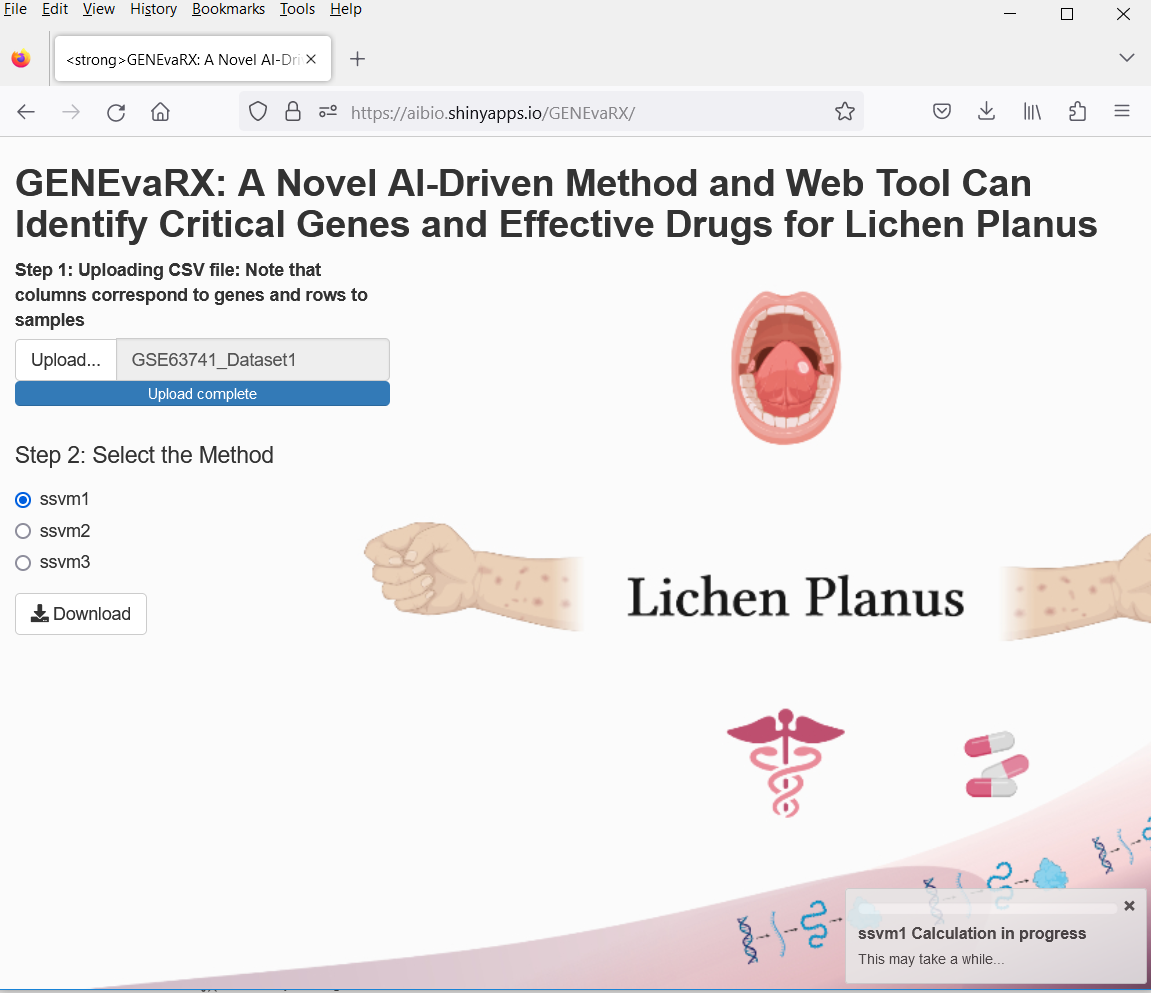

By default, ssvm1 is selected and a progress bar showing the computation status.

After completion, we click on download button to download the results, shown as in the following screen.

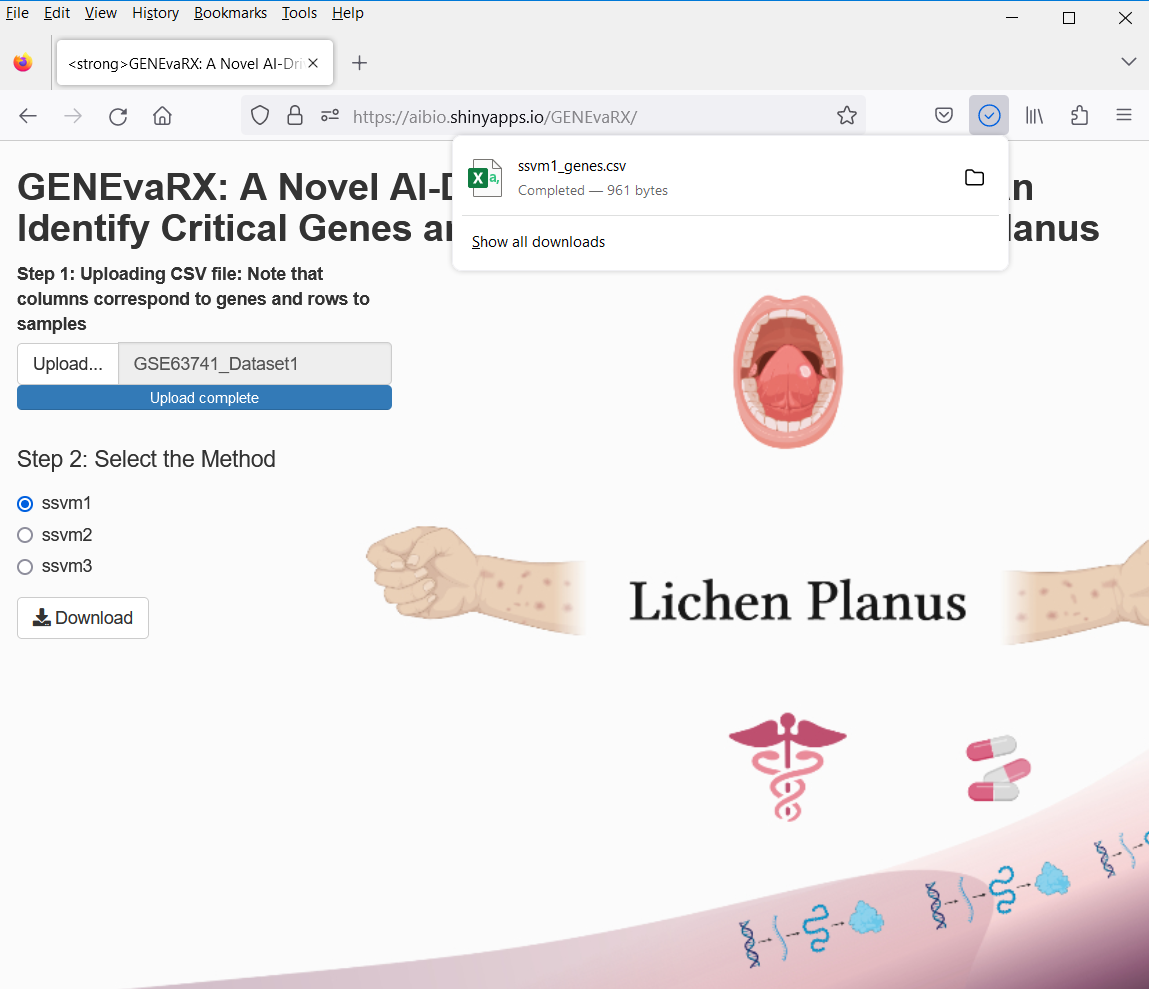

-Now, we open the results in excel file shown as follows

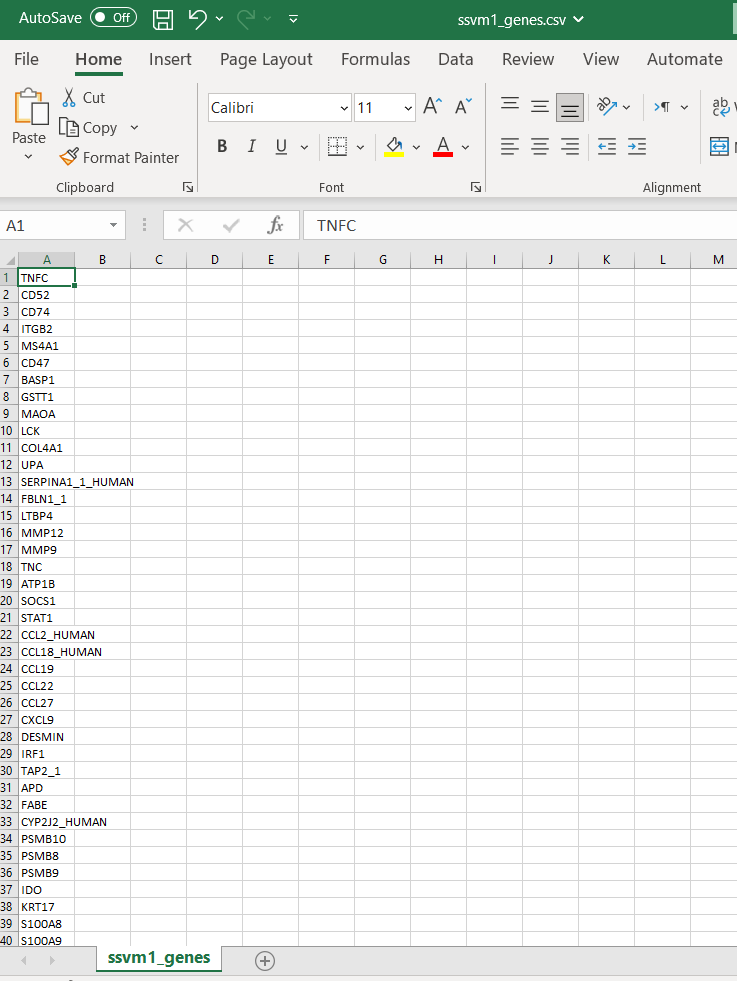

-Suppose we want results of ssvm3, then we select ssvm3 and screen shown as follows

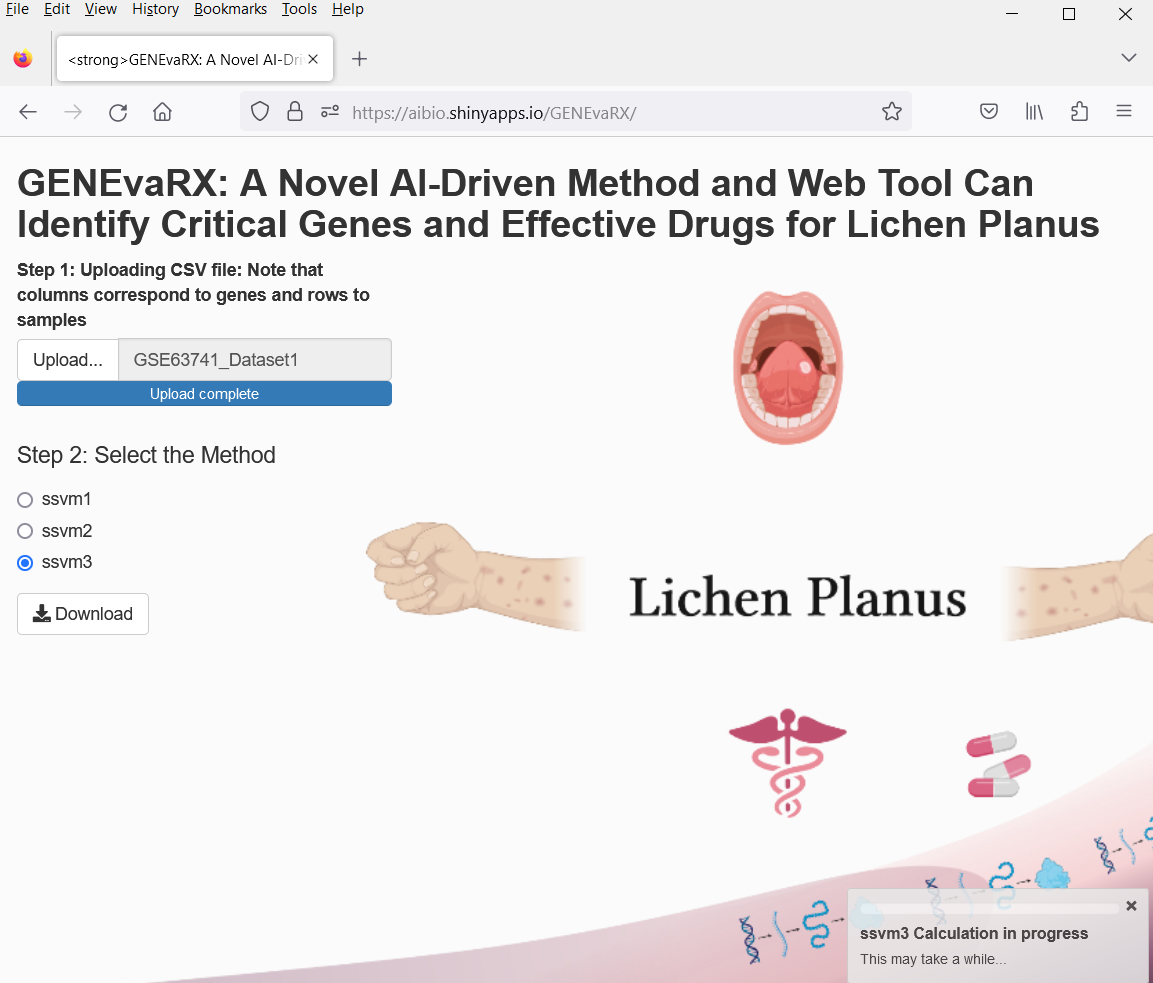

When it completes, we download results as follows.

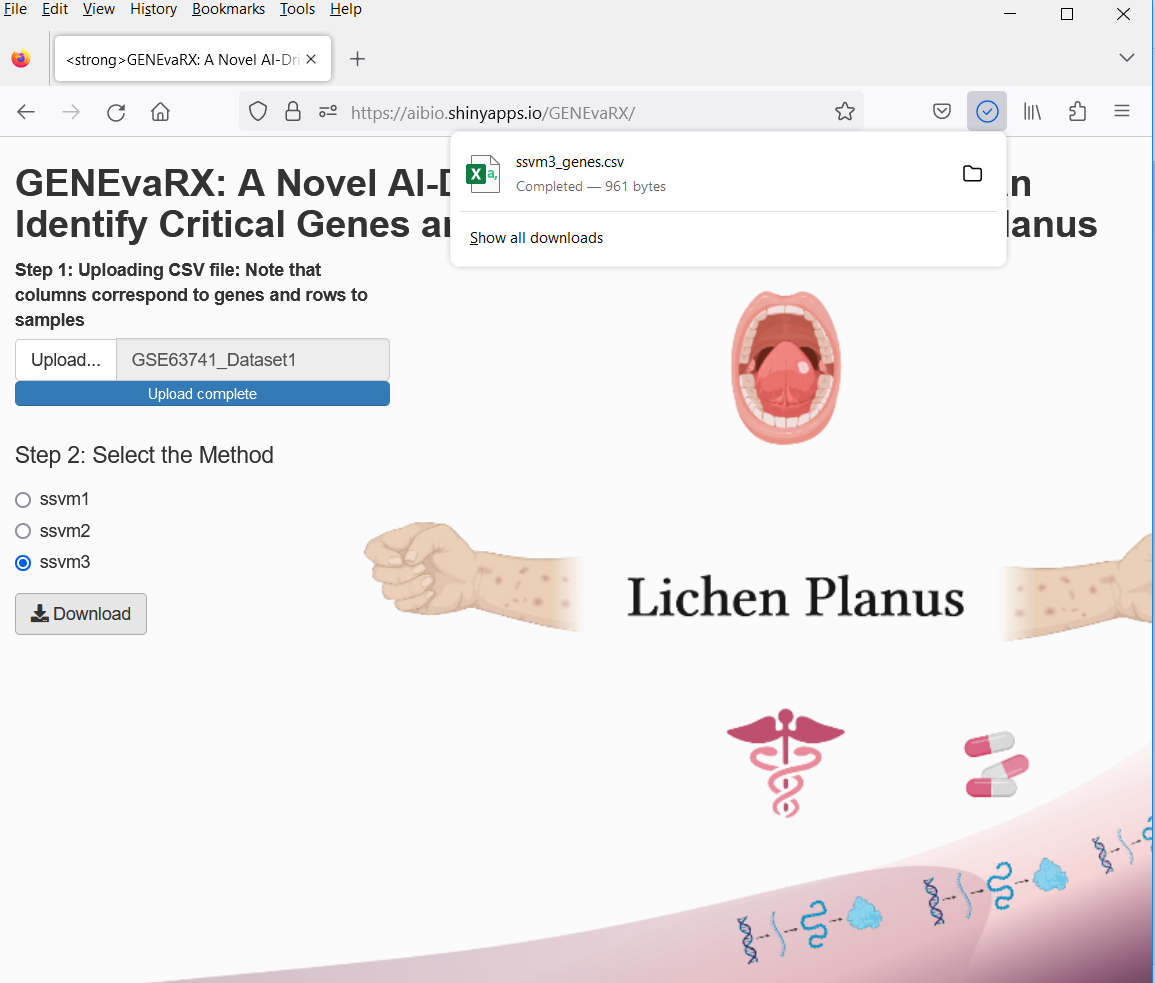

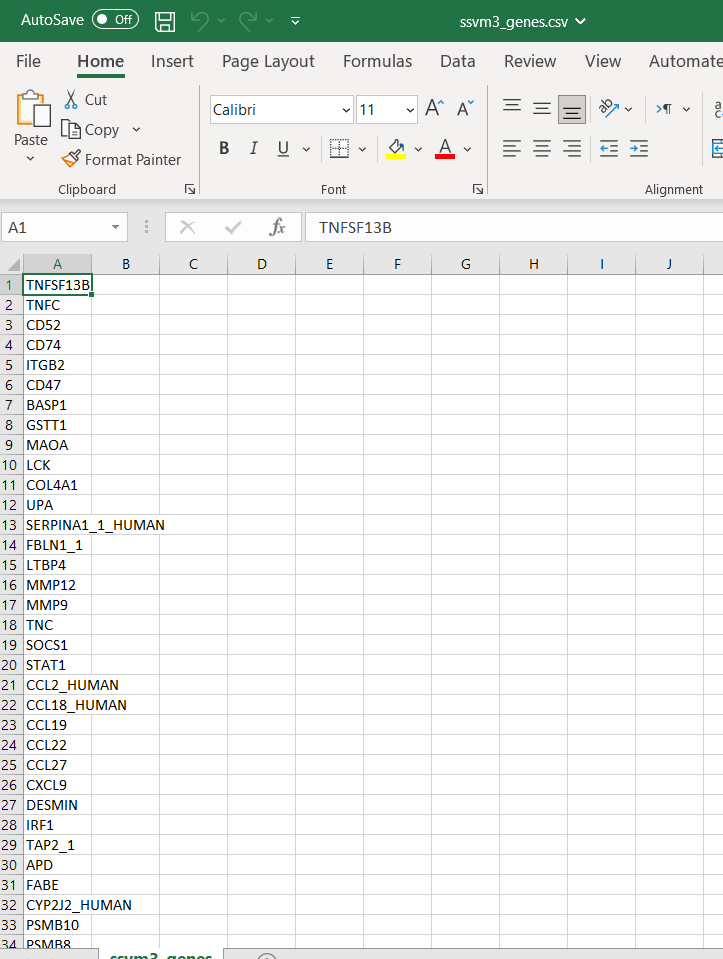
